# Cell-type-specific synaptic scaling mechanisms differentially contribute to associative learning

**DOI:** 10.1101/2025.05.14.654005

**Authors:** Fabio Veneto, Ayça Kepçe, Yue Kris Wu, Julijana Gjorgjieva

## Abstract

Excitatory synaptic scaling regulates network dynamics by proportionally adjusting excitatory synaptic strengths after sensory perturbations. During associative learning, blocking excitatory scaling in conditioned taste aversion paradigms prolongs generalized aversive responses and delays memory specificity. Recent evidence also implicates inhibitory synaptic scaling in the regulation of network dynamics. Specifically, parvalbumin (PV)-expressing inhibitory neurons, targeting perisomatic regions of excitatory (E) pyramidal neurons, and somatostatin (SST)-expressing neurons, targeting distal dendrites, exhibit distinct scaling responses. This leaves open the question of how complex plasticity mechanisms regulate recurrent excitatory-inhibitory circuit dynamics in associative learning. Using computational approaches, we demonstrate that Hebbian plasticity drives memory generalization to novel stimuli not presented during conditioning. Following conditioning, diverse synaptic scaling mechanisms progressively induce memory specificity, which can be regulated by top-down inputs. Our results reveal that, in the absence of excitatory scaling, PV-to-E scaling can effectively compensate and rescue memory specificity, highlighting the presence of degenerate mechanisms in the brain. Notably, in the process of establishing memory specificity, excitatory scaling and PV-to-E scaling function synergistically, while concurrently opposing SST-to-E scaling. The synergistic and antagonistic plasticity mechanisms are orchestrated to shape the temporal evolution of memory representations, from generalized to precise.

**Significance statement:** Associative learning is a fundamental brain function that allows us to link experiences, adapt behavior, and form lasting memories. During this process, memory representations are shaped by synaptic scaling, a homeostatic plasticity mechanism that provides slow, negative feedback to regulate synaptic strengths and adjust network excitability. Operating at the synapses of diverse excitatory and inhibitory cell types, multiple forms of homeostatic plasticity influence the dynamics of associative learning. Here, we demonstrate that synergistic and antagonistic cell-type-specific synaptic scaling mechanisms operate at different types of inhibitory synapses to jointly govern the temporal evolution of memory representations. Through their interaction, they guide the transition from generalized to precise memories.

## Introduction

Synaptic scaling is considered a crucial homeostatic synaptic plasticity mechanism that adjusts the strength of all incoming synapses to a neuron to stabilize network dynamics in response to sensory perturbations (Turrigiano et al., 1998; Turrigiano, 2008; Pozo and Goda, 2010; Tetzlaff, 2011; Wu et al., 2020; Wen and Turrigiano, 2024). Typically studied for synapses between excitatory neurons, this process either downscales or upscales the synapses to compensate for hyperactive or hypoactive activity, respectively (Turrigiano et al., 1998; Kim et al., 2012; Keck et al., 2013; Torrado Pacheco et al., 2021). Unlike Hebbian plasticity, which can occur from seconds to minutes, synaptic scaling operates on a much slower timescale, unfolding over hours to days (Turrigiano et al., 1998; Ibata et al., 2008; Watt, 2010; Keck et al., 2017). Beyond the homeostatic role of excitatory synaptic scaling in compensating for sensory perturbations, recent studies have revealed the importance of excitatory synaptic scaling in associative learning (Wu et al., 2021). Over the years, associative learning has been studied across different sensory modalities (Letzkus et al., 2011; Pakan et al., 2018; Dalmay et al., 2019; Ottenheimer et al., 2023). Recent studies using a classical conditioned taste aversion (CTA) paradigm (Figure 1A), by pairing an aversive unconditioned stimulus (US) with a conditioned stimulus (CS), mice learned the association between the CS and aversion immediately after the conditioning. Four hours after conditioning (early, Figure 1A), mice displayed generalized aversion to a novel test stimulus (TS), a tastant absent during the conditioning. When tested after 24h (intermediate, Figure 1A) and 48h (late, Figure 1A) with the TS, the generalized aversive behavior diminished, and mice exhibited aversion exclusively to the CS, reflecting the formation of memory specificity. Blocking excitatory synaptic scaling significantly prolonged the generalized aversive behavioral response, with mice continuing to exhibit aversion to TS even 24 hours post-conditioning (Figure 1A). Nonetheless, after blocking excitatory synaptic scaling, mice developed memory specificity after 48h post-conditioning, suggesting that other mechanisms may complementarily achieve the specific refinement of memory representations. Here, we sought to understand what these other mechanisms might be.

**Fig. 1.**
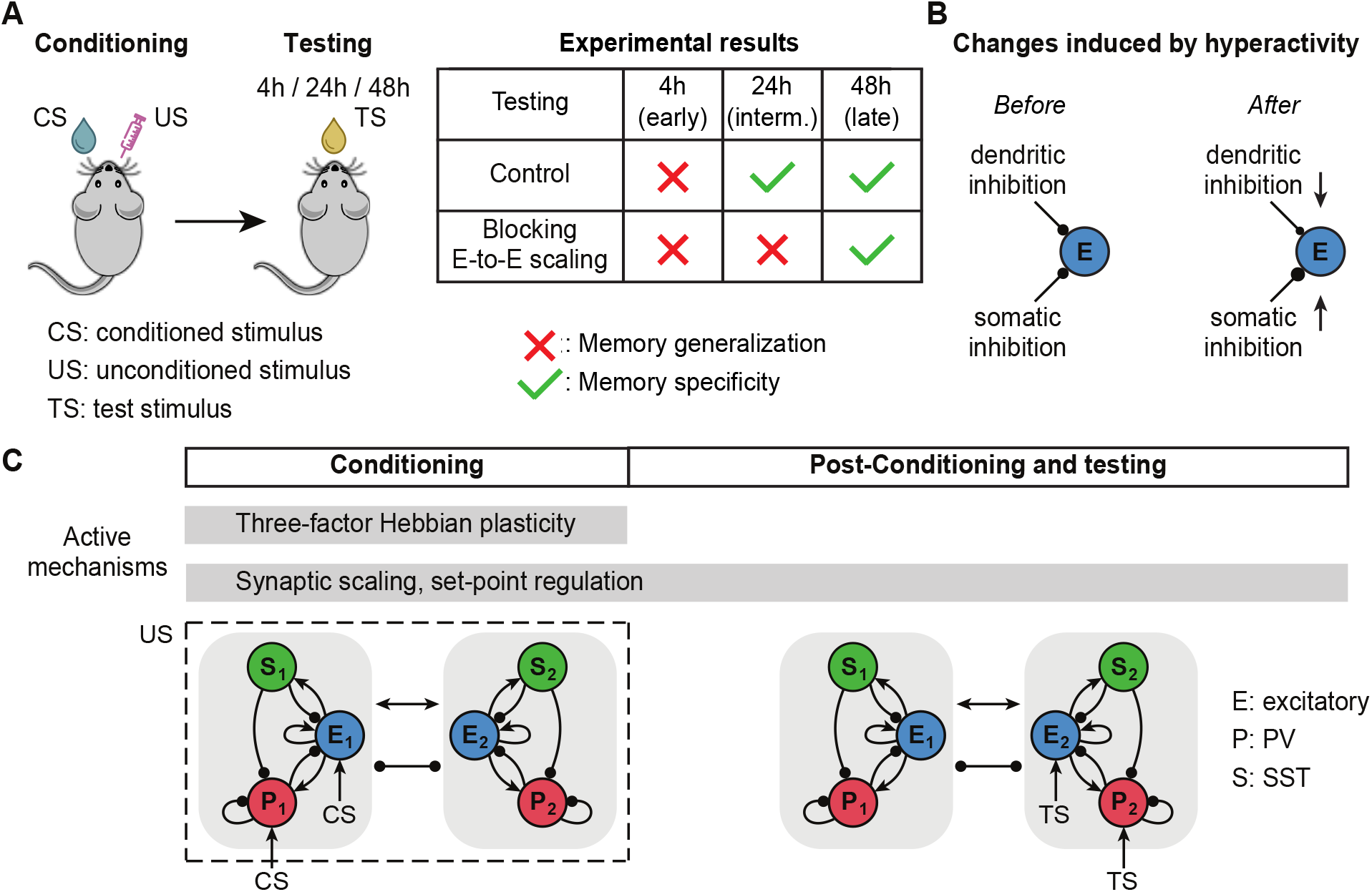
Experimental paradigm and computational framework for associative learning. **A**. Conditioned taste aversion (CTA) paradigm applied in Wu et al. (2021). Conditioning is induced by pairing an aversive unconditioned stimulus (US) with a conditioned stimulus (CS) (left). Memory generalization and specificity are evaluated by measuring the mouse’s aversive behavioral response to a novel test stimulus (TS) at either 4h (early post-conditioning), 24h (intermediate post-conditioning), or 48h (late post-conditioning) (left). Mice exhibit an aversive behavioral response to TS at 4h but not at 24h and 48h, indicating a switching from memory generalization to memory specificity (right). When blocking excitatory scaling, memory generalization persists at 24h but diminishes by 48h (right). **B**. Target-specific inhibitory synaptic scaling reported in Prestigio et al. (2021). Hyperactivity in postsynaptic excitatory neurons induces a downscaling of dendritic inhibition while upscaling somatic inhibition. **C**. Schematic of network model with two subnetworks. Each subnetwork consists of one excitatory, one PV and one SST population. Different subnetworks are tuned to different stimuli corresponding to different tastants in the conditioned taste aversion experiments. During conditioning, excitatory (E1) and PV (P1) populations in subnetwork 1 receive additional inputs corresponding to a CS, while the US is present. During the test period, excitatory (E2) and PV (P2) populations in subnetwork 2 receive additional inputs corresponding to a TS. Three-factor Hebbian plasticity operates during conditioning, whereas synaptic scaling and set point regulation mechanisms are active during both conditioning and post-conditioning phases.

In addition to excitatory synaptic scaling, inhibitory synaptic scaling has also been found to regulate network dynamics in response to sensory perturbations (Kilman et al., 2002; Swanwick et al., 2006; Prestigio et al., 2021). Notably, the scaling of inhibitory synapses is target-dependent (Prestigio et al., 2021), whereby hyperactivity in excitatory pyramidal neurons (E) induces upscaling of inhibitory synapses at perisomatic regions of excitatory neurons while unexpectedly downscaling of inhibitory synapses at dendritic regions. These connectivity preferences have been related to molecularly distinct interneuron subtypes (Tremblay et al., 2016) (Figure 1B). For instance, parvalbumin (PV)-expressing inhibitory neurons primarily innervate perisomatic regions, whereas somatostatin (SST)-expressing inhibitory neurons predominantly target distal dendritic regions (Lazarus and Huang, 2011; Hioki et al., 2013; Dorsett et al., 2021). Yet, how these inhibitory synaptic scaling mechanisms interact with the well-established excitatory Hebbian and synaptic scaling mechanisms during associative learning remains elusive.

In this work, we combined analytical calculations and numerical simulations to demonstrate that rapid Hebbian plasticity drives memory generalization to novel test stimuli that are absent during conditioning. Following conditioning, we find that different forms of synaptic scaling regulate neural dynamics, progressively inducing memory specificity over time. Our findings highlight the critical role of different forms of synaptic scaling in achieving precise memory representations and propose a role for top-down inputs in modulating associative learning. When excitatory scaling is absent, memory specificity can be rescued by PV-to-E scaling, indicating the existence of degenerate mechanisms in the brain. We find that excitatory scaling and PV-to-E scaling work synergistically while counteracting the effects of SST-to-E scaling. This intricate interplay between synergistic and antagonistic plasticity mechanisms drives the temporal evolution of memory specificity, facilitating a smooth transition from generalized to specific memory representations.

## Results

To investigate how different plasticity mechanisms – rapid Hebbian and slower forms of synaptic scaling – interact with each other and give rise to memory specificity in associative learning, we developed a rate-based recurrent network model consisting of two interconnected subnetworks. Each subnetwork includes one excitatory (E) population and two distinct inhibitory populations: PV and SST (see Methods, Figure 1C). We assume that different subnetworks are tuned to different stimuli corresponding to different tastants in the conditioned taste aversion experiments. Inspired by experimental studies indicating that PV inhibitory neurons primarily innervate perisomatic regions, while SST inhibitory neurons predominantly target distal dendritic regions, we modeled somatic inhibition to the excitatory population as coming from the PV population and dendritic inhibition as coming from the SST population (Figure 1C) (Lazarus and Huang, 2011; Pfeffer et al., 2013; Dorsett et al., 2021). The network connectivity was designed to incorporate previously reported experimental features, including the absence of inhibitory connections from PV and SST interneurons to SST interneurons (Pfeffer et al., 2013).

### Three-factor Hebbian plasticity strengthens excitatory-to-excitatory connections during conditioning

To model the conditioning procedure in the conditioned taste aversion paradigm, the excitatory (E_1_) and PV (PV_1_) populations in subnetwork 1 receive additional inputs to represent the conditioned stimulus (CS) (Figure 2A) (Ji et al., 2015). Inspired by experimental studies demonstrating that reward or punishment plays a crucial role in learning (Pawlak, 2010; Gerstner et al., 2018), we applied a three-factor Hebbian learning rule to update the E-to-E connection strength during conditioning:

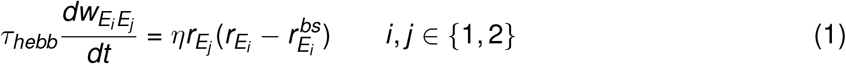

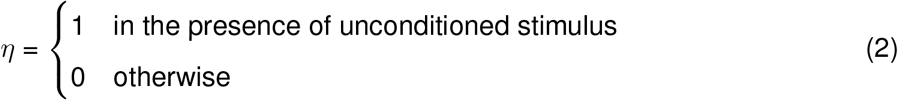

where *τ*_*hebb*_ is the time constant of Hebbian plasticity, 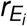 denotes the activity of the excitatory population in subnetwork *i* with the superscript ‘bs’ representing the baseline activity before conditioning, *i, j* representing the indices of subnetworks. The presence of the aversive unconditioned stimulus (US) determines the third factor *η* and serves as a gate for Hebbian plasticity, enabling plasticity during the conditioning phase while disabling it elsewhere.

**Fig. 2.**
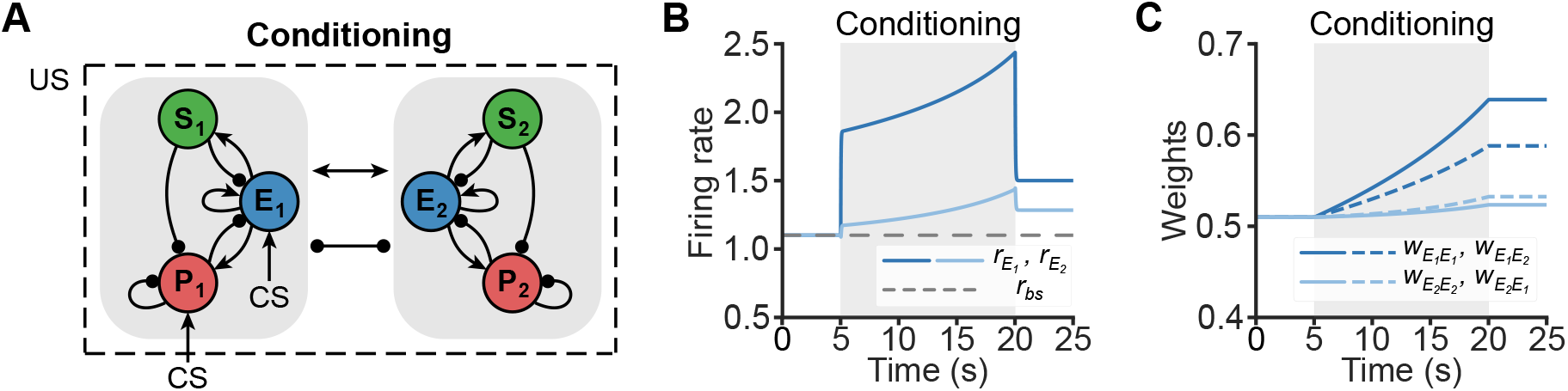
Hebbian plasticity enhances excitatory activity and strengthens E-to-E connections during conditioning. **A**. Network schematic of the conditioning phase. During conditioning, E and PV populations of subnetwork 1 receive additional inputs that correspond to the conditioned stimulus. During this phase, the unconditioned stimulus is present, modulating Hebbian plasticity. **B**. Activity of excitatory population in subnetwork 1 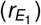 and subnetwork 2 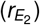. Conditioning is applied during the interval from 5 to 20s by increasing the inputs to E and PV populations in subnetwork 1. The dashed line represents the baseline activity level measured before conditioning. **C**. Excitatory to excitatory connection strength during conditioning. Different connections are indicated by the differently colored lines.

During the simulation of conditioning, the CS leads to an increase in the excitatory activity in sub-network 1 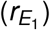 (Figure 2B). Despite not being directly stimulated by the CS, the excitatory activity in subnetwork 2 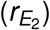 also increases, albeit to a lesser extent, due to recurrent excitatory connections between *E*_1_ and *E*_2_ (Figure 2B). During conditioning, in the presence of the US, E-to-E connection strengths increase through Hebbian plasticity, with the strongest enhancement observed in the connection strength within the excitatory population of subnetwork 1 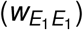 (Figure 2C).

### Memory undergoes transient generalization caused by Hebbian plasticity before gradually achieving specificity

Together with Hebbian plasticity acting on excitatory-to-excitatory synapses during conditioning, we incorporated synaptic scaling at the connections from E-to-E synapses, i.e., excitatory scaling. This is consistent with experimental findings that hyperactivity (hypoactivity) of excitatory neurons leads to downscaling (upscaling) of E-to-E synapses. Synaptic scaling adjusts synaptic weights to maintain stable activity levels, preventing activity from becoming excessively low or high (Turrigiano et al., 1998; Kim et al., 2012; Keck et al., 2013; Torrado Pacheco et al., 2021). This process is generally considered to be multiplicative and independent of presynaptic activity (Turrigiano, 2008). Following previous computational studies (Van Rossum et al., 2000), we modeled the change of connection strength from the excitatory population in subnetwork *j* to the excitatory population in subnetwork *i* via synaptic scaling as follows:

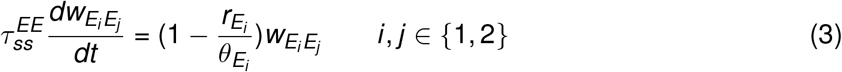

Here, *τ*_*ss*_ represents the time constant of synaptic scaling for individual type of connections,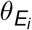 denotes the target firing rate or the set point of the excitatory population in the subnetwork *i*.

To investigate how different inhibitory synaptic scaling, discovered experimentally to excitatory dendrites and somas (Prestigio et al., 2021), collectively affect associative learning, we implemented synaptic scaling from PV-to-E and SST-to-E synapses. In line with the observed decrease in somatic inhibition induced by the hyperactivity of postsynaptic excitatory neurons (Prestigio et al., 2021), PV-to-E synapses are scaled by:

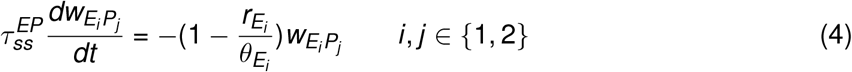

Similarly, consistent with the observed increase in dendritic inhibition resulting from the hyperactivity of postsynaptic excitatory neurons (Prestigio et al., 2021), SST-to-E synapses are scaled by:

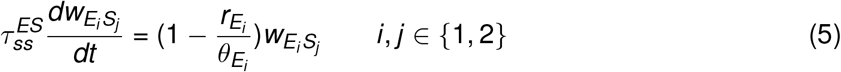

The set points of the two excitatory populations were allowed to change (Leman et al., 2025), to reflect distinct dynamics and activity levels that emerge due to direct stimulation of one population during the conditioning phase. In particular, the set points were jointly determined by the corresponding activity and the set point regulator *β* according to the following dynamics:

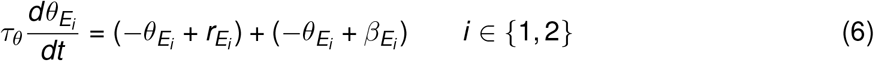

where *τ*_*θ*_ is the time constant governing the plasticity of the set points. The set point regulator *β* can be considered a form of a network homeostatic mechanism that uniformly regulates the activity of the entire network and is dynamically updated according to:

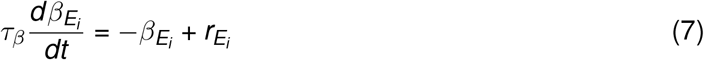

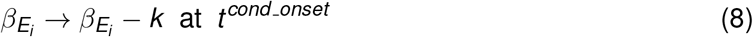

where *τ*_*β*_ denotes the time constant governing the plasticity of the set point regulator and *k* is a free parameter that determines the magnitude of the abrupt decrease in the set point regulator *β* of excitatory populations in both subnetworks at the onset of conditioning *t*^*cond_onset*^ . Conditioning raises the excitatory activity *r*_*E*_, thereby increasing both the set point *θ* and the set point regulator *β*. In contrast, this sudden reduction in *β* counteracts the increases induced by conditioning and functions as a homeostatic mechanism to globally regulate the overall activity level. While it might be possible to directly incorporate the parameter *k* into the set point dynamics, from a biological perspective a slowly changing set point regulator is more consistent with activity-dependent gene expression processes linked to slow homeostatic plasticity. The abrupt change in the set point regulator at condition onset was implemented for modeling convenience. Biologically, such a change of set point regulator might be more naturally associated with the presence of the unconditioned stimulus (e.g. reward or punishment) rather than a precise onset time point.

To evaluate memory specificity after conditioning, we presented a test stimulus (TS) to the network by providing excitatory inputs to the E and PV populations in subnetwork 2 at three distinct test time points (4h, 24h, and 48h), and measured the activity of the excitatory population in subnetwork 1, denoted by 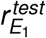 (Figure 3A). This activity is compared to a reference activity, 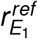, obtained by simulating the network under identical initial conditions but without applying conditioning (Figure S1). We observed that 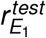 exceeds 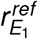 during TS presentation at 4h (Figure 3B), whereas, at 24h and 48h, 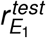 is smaller than 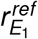 (Figure 3B). These results suggest that, following conditioning, the memory initially generalizes to test stimuli but eventually becomes specific over time.

**Fig. 3.**
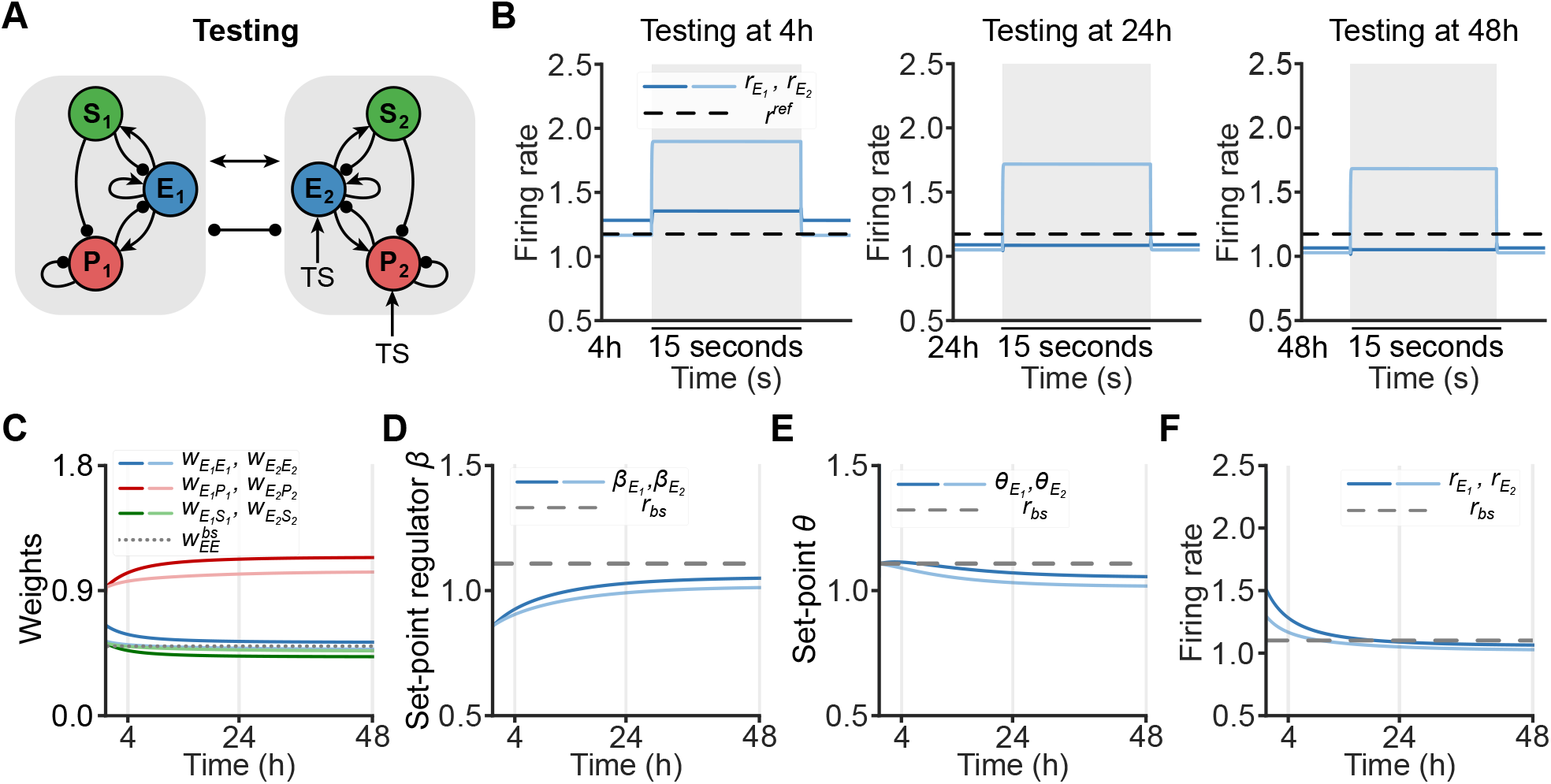
Memory gradually transitions from generalization to specificity. **A**. Network schematic of the testing phase. After conditioning, E (E2) and PV (P2) populations of subnetwork 2 receive additional inputs that correspond to the test stimulus. The unconditioned stimulus is not presented during this phase. **B**. Responses of excitatory populations in subnetwork 1 and subnetwork 2 when presenting a test stimulus for 15s (gray) at 4h (left), 24h (middle) and 48h (right). The black horizontal lines indicate the reference activity (*r* ^*ref*^), measured by the excitatory population in subnetwork 1 in response to a test stimulus under identical initial weights conditions but without plasticity. Here, a moderate value of *k* = 0.25 is applied. **C**. Different connection strengths (E-to-E, PV-to-E and SST-to-E) after conditioning. The horizontal line indicates the baseline excitatory to excitatory connection strength 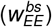 before conditioning. **D**. Evolution of set points regulators of excitatory population in subnetwork 1 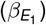 and subnetwork 2 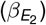 after conditioning up to 48h. The gray horizontal dashed line represents the baseline activity level measured before conditioning. **E**. Same as D but for set point *θ*. **F**. Activity of excitatory population in subnetwork 1 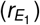 and subnetwork 2 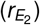 after conditioning.

Although synaptic scaling is active throughout the entire simulation, including the conditioning phase, its slow time constant renders changes in the E-to-E connection strength during conditioning negligible. Following conditioning, the weights evolve solely due to the different forms of synaptic scaling, undergoing significant changes over time: E-to-E weights decrease, PV-to-E weights increase, and SST-to-E weights decrease (Figure 3C). Note that despite the decrease in E-to-E weights after conditioning, the E-to-E weight in subnetwork 1 remains above the preconditioning level (Figure 3C), consistent with the idea that associative memory formation involves the strengthening of synaptic weights. Following the abrupt decrease at the onset of conditioning, the set point regulators gradually rise over time, driven by increased excitatory activity in the early post-conditioning period (Figure 3D). Due to the set point regulators being lower than the baseline activity, the set points, decreased throughout the post-conditioning period, eventually stabilizing at a new steady state lower than the initial set points (Figure 3E). Consequently, following conditioning, the excitatory activity, along with PV and SST activity, gradually decreases over time and converges towards the new set points (Figure 3F). Our results indicate that synaptic scaling gradually reshapes network connectivity after conditioning, driving a progressive reduction in excitatory, PV, and SST activity (Figure S2) as the system stabilizes to a new equilibrium.

### Characterization of the temporal evolution of memory representations

To characterize how memory representations dynamically evolve over time, we aimed to describe the network’s response to the test stimulus following conditioning in the model. To that end, we introduced a procedure consisting of two phases for calculating the weights in the network during and post-conditioning, followed by the testing phase where we defined a “Generalization Index” to measure the degree of memory specificity or generalization (Figure 4A).

**Fig. 4.**
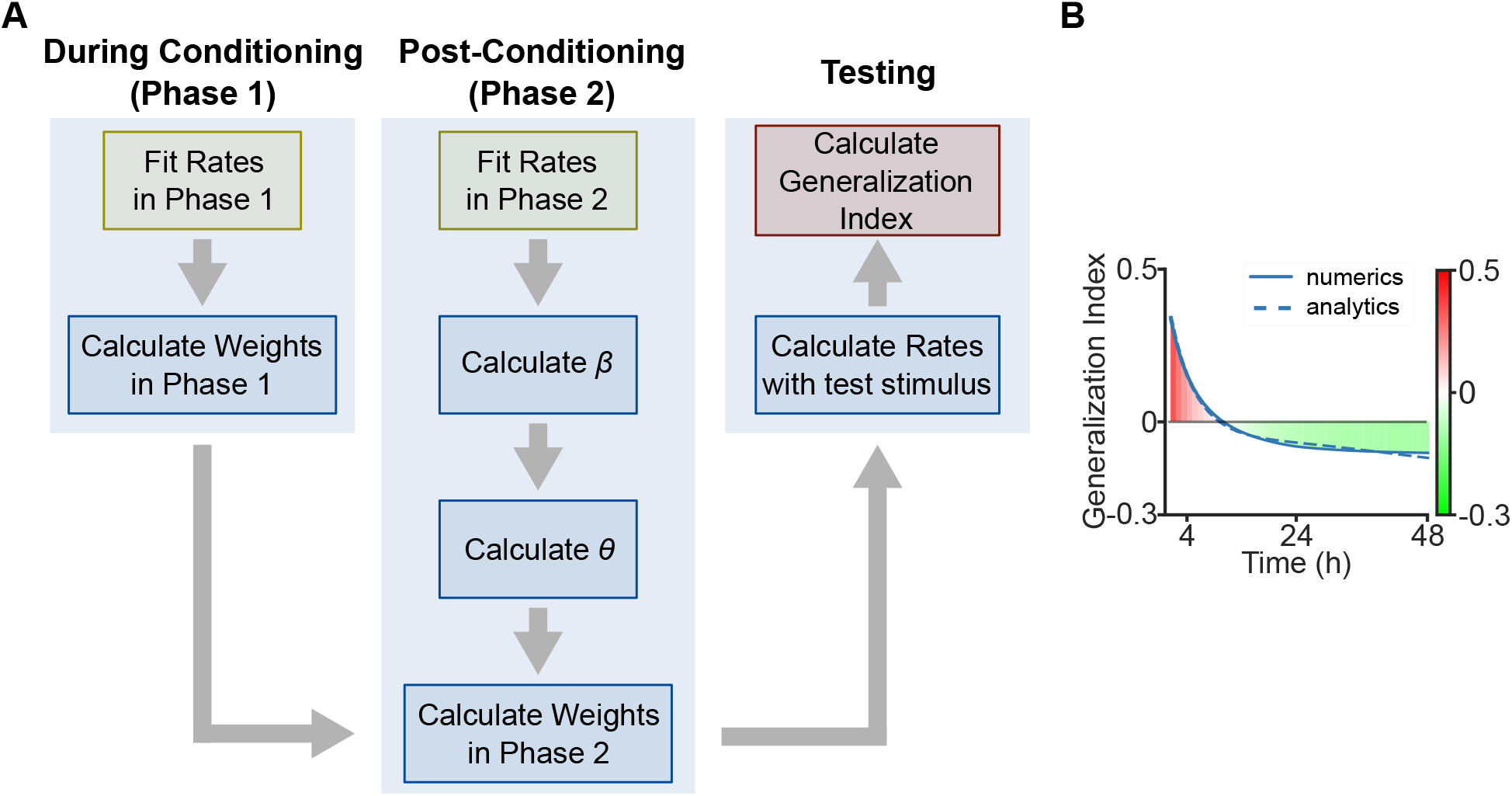
A procedure to assess the evolution of memory representations. **A**. Workflow chart for calculating the Generalization Index (GI) (Eq. 9), see main text. **B**. Evolution of the GI after conditioning. Numerical results (solid line) represent GI measurements taken hourly post-conditioning, while analytical results are derived from continuous GI calculations using the procedure described in Figure 4A. The GI shifts from positive to negative, indicating the transition from memory generalization to memory specificity.

During conditioning (Phase 1), the excitatory firing rate can be well approximated by an exponential function (see Methods), capturing the simulated activity dynamics (Figure S3). We derived the evolution of E-to-E synaptic weights during this phase from solving the dynamics of the three-factor Hebbian learning rule, producing values that closely match those observed in simulations (Figure S3). Given that synaptic scaling operates on a substantially longer timescale than the duration of conditioning, its effects are negligible in this phase. After conditioning (Phase 2), excitatory firing rates can also be well described by an exponential function allowing us to compute the set point regulator *β*. Subsequently, we derived the set point *θ* from the obtained *β*. This allows us to accurately determine the synaptic weight evolution during post-conditioning (Figure S3).

To quantify the degree of memory specificity or generalization, we defined a new measure, called the Generalization Index (GI):

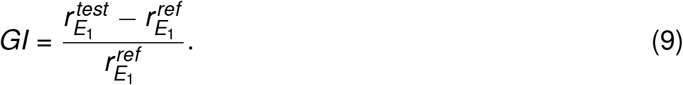

The GI quantifies the relative change in the excitatory population activity of subnetwork 1 between the test and the previously defined reference conditions. A positive GI (e.g. 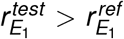) suggests that a memory has been generalized, a negative GI (e.g. 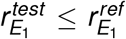) indicates that a memory is specific. The magnitude of GI reflects the strength of memory specificity or generalization. Applying the above procedure, we found that the GI gradually transitions from positive to negative (Figure 4B). This transition suggests that, following conditioning, the memory initially generalizes to test stimuli but gradually becomes specific over time. Taken together, our procedure provides a quantitative characterization of the temporal dynamics underlying evolving memory representations, revealing how these representations are progressively reshaped over time. Furthermore, as chronic neural activity recordings become increasingly available, the analytical procedure could serve as a useful predictive tool for assessing memory generalization over time.

### The network homeostatic mechanism adjusts the set points and the strength of memory specificity

The network homeostatic mechanism in our model (Eq. 8), governed by the parameter *k*, influences the set point regulators and thus the new set points. For a small *k*, in the absence of the network homeostatic mechanism, the set points *θ* slightly increase throughout the post-conditioning period and stabilize at a new steady state moderately above the baseline activity (i.e., the initial set points) (Figure 5A). Following conditioning, excitatory activity decreases and approaches the new set points (Figure 5B). The GI remains positive throughout the post-conditioning period consistent with memory generalization (Figure 5C). In contrast, increasing the influence of the network homeostatic mechanism (large *k*), suppresses the set points *θ* and excitatory activity (Figure 5D, E). In this case, the GI shifts from positive to negative throughout the post-conditioning period (Figure 5F) and reaches a lower value compared to the immediate *k* condition (Figure 3C), indicating enhanced memory specificity. Taken together, these results suggest that the network homeostatic mechanism significantly influences the set points and regulates the degree of memory specificity.

**Fig. 5.**
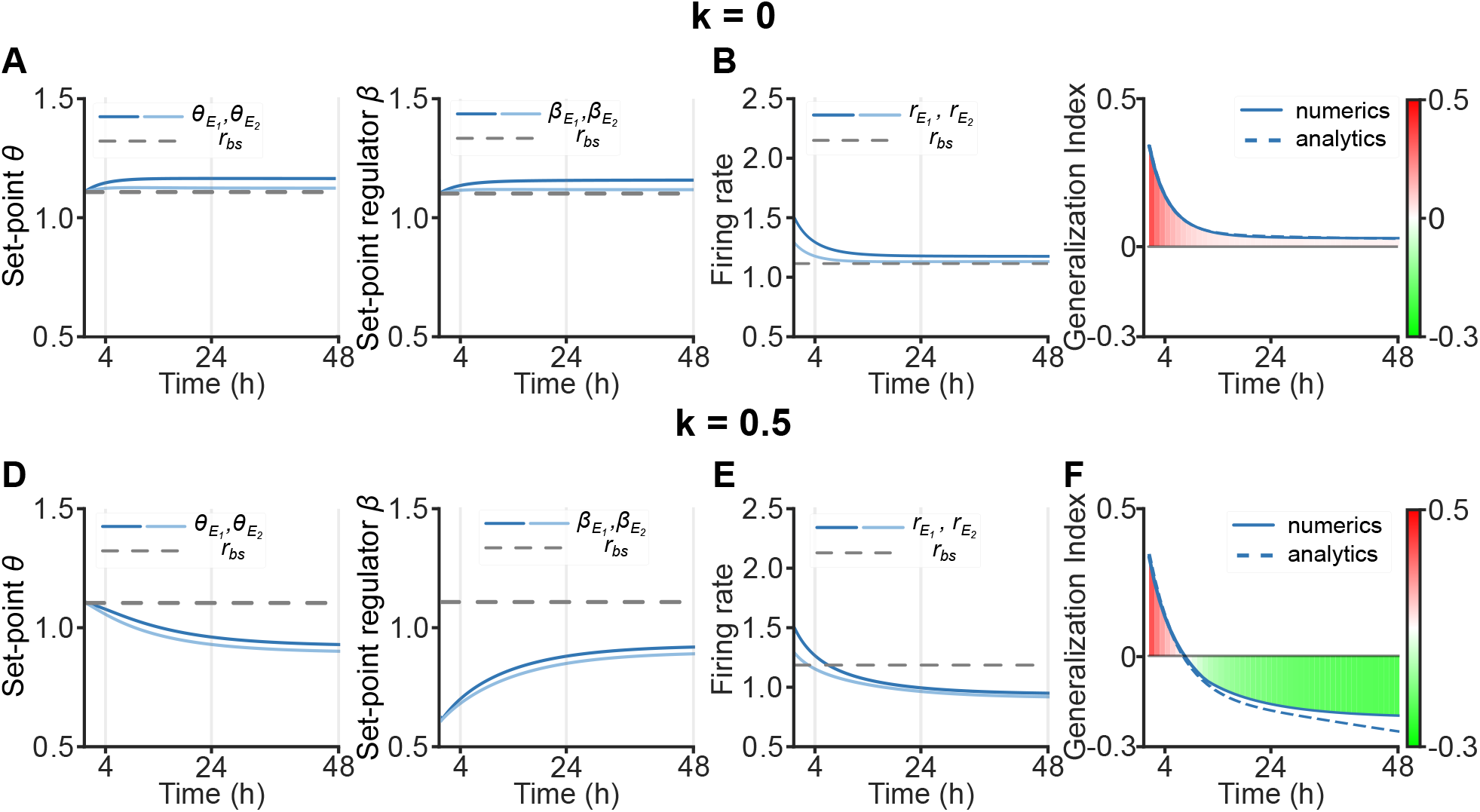
The network homeostatic mechanism regulates the emergence of memory specificity. **A**. Evolution of set point *θ* (left) and set point regulator *β* (right) in subnetwork 1 and subnetwork 2 after conditioning up to 48h for *k* = 0, corresponding to the absence of the network homeostatic mechanism. The gray horizontal dashed line represents the baseline activity level measured before conditioning. **B**. Evolution of excitatory firing rate in subnetwork 1 and subnetwork 2 after conditioning up to 48h for *k* = 0. **C**. Evolution of the GI after conditioning. For *k* = 0, GI remains positive after conditioning. A positive GI indicates memory generalization, whereas a negative GI represents memory specificity. The dashed line indicates the analytically derived GI. **D - F**. Same as (A - C) but for *k* = 0.5. The GI transitions from positive to negative after 4h after conditioning.

### Top-down inputs regulate memory specificity

In addition to bottom-up inputs driven by sensory stimuli, primary sensory cortical areas also receive abundant top-down inputs from higher-order regions which influence neuronal processing in local recurrent circuits (Johnson and Burkhalter, 1997; Garrett et al., 2014). To investigate how top-down inputs influence associative learning, we applied an additional input to SST populations (a common target of top-down inputs) during conditioning (Batista-Brito et al., 2018; Shen et al., 2022). When introducing an inhibitory top-down input to both SST populations (Figure 6A), we found that inhibition of SST interneurons disinhibits excitatory neurons, leading to a drastic increase in the firing rates of both subnetworks during conditioning (Figure 6B). However, after conditioning, excitatory activity rapidly declines (Figure 6C). Furthermore, due to the large initial change in firing rates, the synaptic weights undergo substantial modification (Figure 6D). The GI transitions from positive to negative later in the post-conditioning period than in the absence of inhibitory top-down input, indicating a slower emergence of memory specificity (Figure 6E). In contrast, when applying excitatory top-down input to both SST populations in the same amount during conditioning (Figure 6F), the excitatory firing rate of subnetwork 1 slightly increases, while subnetwork 2 decreases (Figure 6G). After conditioning, both subnetworks’ excitatory activity gradually decline below baseline levels (Figure 6H). The small deviation of excitatory activity from baseline results in minor changes in inhibitory synaptic weights (Figure 6I). The GI transitions from positive to negative earlier in the post-conditioning period than in the absence of inhibitory top-down input, indicating a faster emergence of memory specificity (Figure 6J).

**Fig. 6.**
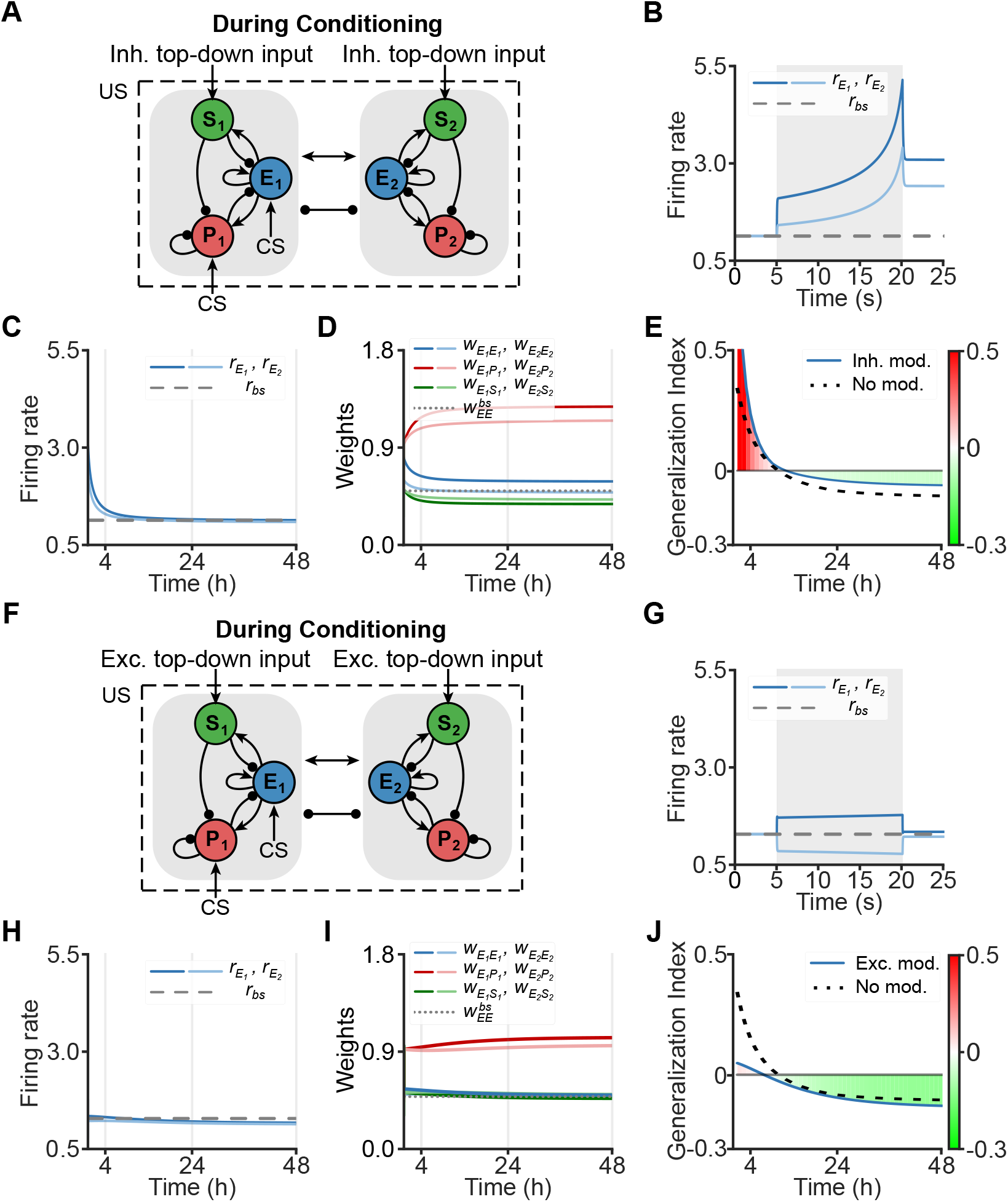
Top-down inputs influence memory specificity. **A**. Network schematic of the conditioning phase in the presence of inhibitory top-down inputs. During conditioning, the E and PV populations of subnetwork 1 receive additional inputs that correspond to the conditioned stimulus, while the SST population of both subnetworks 1 and 2 receives additional inhibitory top-down inputs. **B**. Activity of excitatory population in subnetwork 1 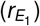 and subnetwork 2 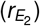 during conditioning in the presence of inhibitory top-down inputs. Conditioning is marked by the gray interval from 5 to 20s. The gray horizontal dashed line represents the baseline activity level measured before conditioning. **C**. Activity of excitatory population in subnetwork 1 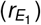 and subnetwork 2 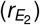 after conditioning in the presence of inhibitory top-down inputs. **D**. Different connection strengths (E-to-E, PV-to-E and SST-to-E) after conditioning. **E**. Evolution of the Generalization Index (GI) after conditioning in the presence (blue solid) and in the absence (black dashed) of inhibitory top-down inputs. A positive GI indicates memory generalization, whereas a negative GI represents memory specificity. **F - J**. Same as (A - E) but in the presence of excitatory top-down inputs.

These findings suggest that top-down inhibition of SST would transiently enhance excitatory activity and slow memory specificity, whereas top-down excitation of SST confines the degree of increase in excitatory activity and accelerates the refinement of memory representations.

### Synaptic scaling is essential for associative learning, with distinct contributions from specific cell types

Next, we investigated the role of synaptic scaling in associative learning by blocking all synaptic scaling mechanisms. In the absence of synaptic scaling mechanisms, both firing rates and weights stay unchanged during the post-conditioning period (Figure 7A), 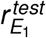 exceeded *r* ^*ref*^ (Figure 7B), and the GI remains positive (Figure 7C), suggesting memory generalization. Together, these results indicate that synaptic scaling is crucial for achieving memory specificity.

**Fig. 7.**
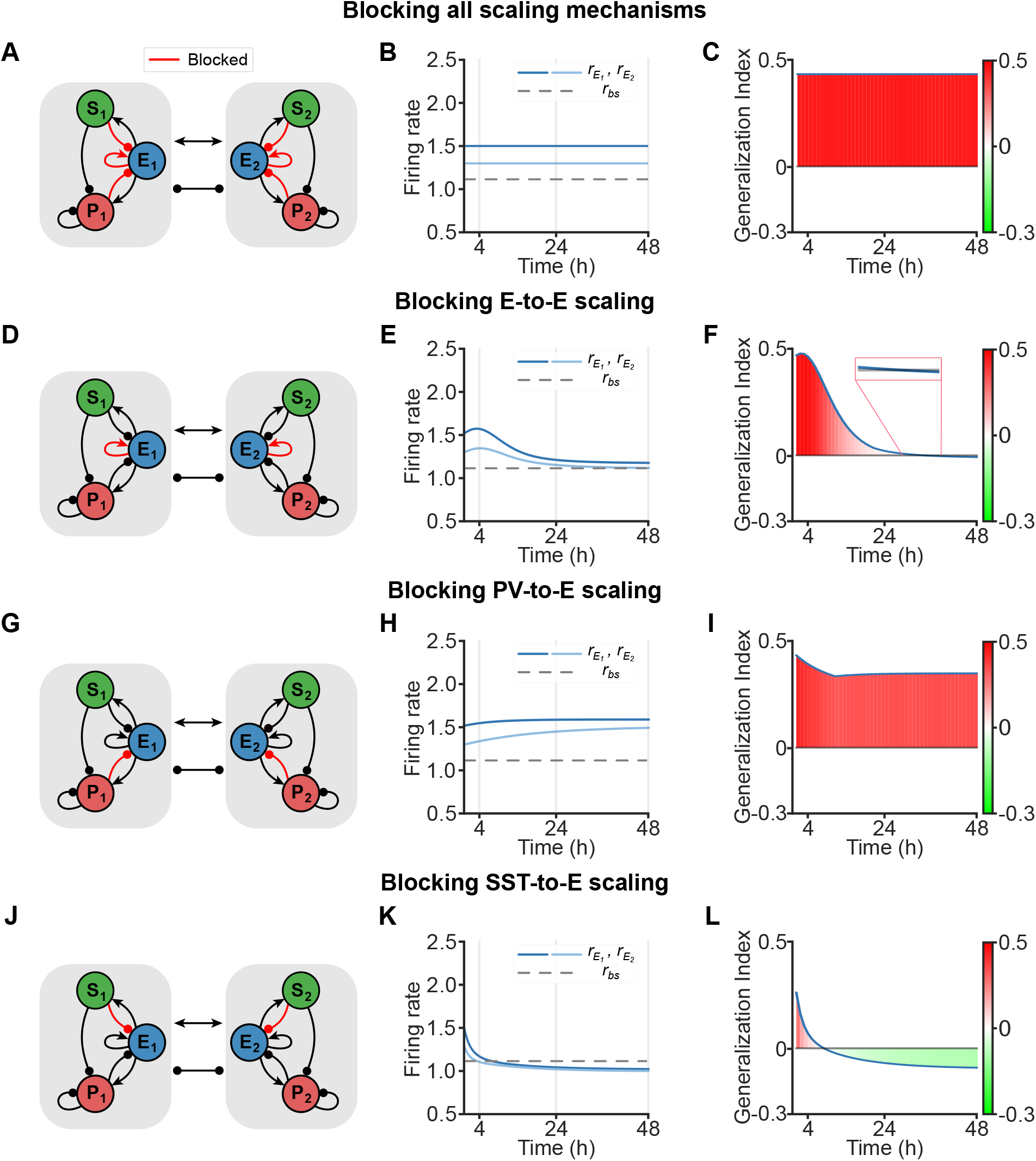
Cell-type-specific synaptic scaling contributions to memory refinement in associative learning. **A**. Network schematic when blocking all scaling mechanisms (red connections). Cross-connections are also blocked accordingly. **B**. Activity of excitatory population in subnetwork 1 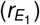 and subnetwork 2 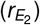 after conditioning when blocking all scaling mechanisms. The gray horizontal dashed line represents the baseline activity level measured before conditioning. **C**. Evolution of the Generalization Index (GI) after conditioning when blocking all scaling mechanisms. A positive GI indicates memory generalization, whereas a negative GI represents memory specificity. **D - F**. Same as (A - C) but for blocking E-to-E scaling. **G - I**. Same as (A - C) but for blocking PV-to-E scaling. **J - L**. Same as (A - C) but for blocking SST-to-E scaling.

But to which extent do the different types of synaptic scaling affect associative learning? When selectively blocking E-to-E scaling (Figure 7D), we found that excitatory firing rates of both subnetworks gradually converge to levels close to their baseline (Figure 7E). The GI shifts from positive to negative (Figure 7F), albeit at a later time point compared to when all scaling mechanisms are present (Figure 4B) indicating that the memory eventually becomes specific.

Blocking PV-to-E scaling (Figure 7G) elevates excitatory firing rates of both subnetworks constantly beyond their baseline levels (Figure 7H), resulting in a positive GI and hence generalized memories (Figure 7I). In contrast, when SST-to-E scaling is blocked (Figure 7J), the excitatory firing rates of both subnetworks promptly decrease to levels below their baseline (Figure 7K). This results in the GI transitioning from positive to negative earlier in the post-conditioning period than with intact scaling mechanisms (Figure 7L), indicating a more rapid emergence of memory specificity. To assess the robustness of the observed results to parameter selection, we conducted numerous simulations using different initial weight conditions (see Methods). We found that, in 93% of initial weight conditions, PV-to-E scaling is essential to achieve memory specificity, whereas blocking SST-to-E scaling always accelerates the transition to memory specificity (see Methods, Figure S4, S5). Together, these results suggest that E-to-E and PV-to-E scaling operate synergistically, while SST-to-E scaling acts antagonistically, to collectively regulate the timing of memory specificity.

### Multi-compartment model confirms different contributions of cell-type-specific synaptic scaling to associative learning

To assess whether our main conclusions from the simplified point neuron model also hold in a more biologically realistic setting, we implemented a multi-compartment model of excitatory neurons consisting of three compartments: soma, basal dendrite, and apical dendrite, whereas two distinct groups of inhibitory neurons are modeled as point neurons targeting the soma and dendrites (both basal and apical dendrites) of the excitatory populations, respectively (Figure 8A, see Methods). In contrast to the point neuron model, in this multi-compartment model, the scaling of the inhibitory neurons resembling PV acts exclusively on the soma of the excitatory populations, whereas the scaling of the other inhibitory neurons resembling SST operates solely on the basal and apical dendrites. This setup enables a direct correspondence to experimental findings on target-dependent inhibitory synaptic scaling (Prestigio et al., 2021).

**Fig. 8.**
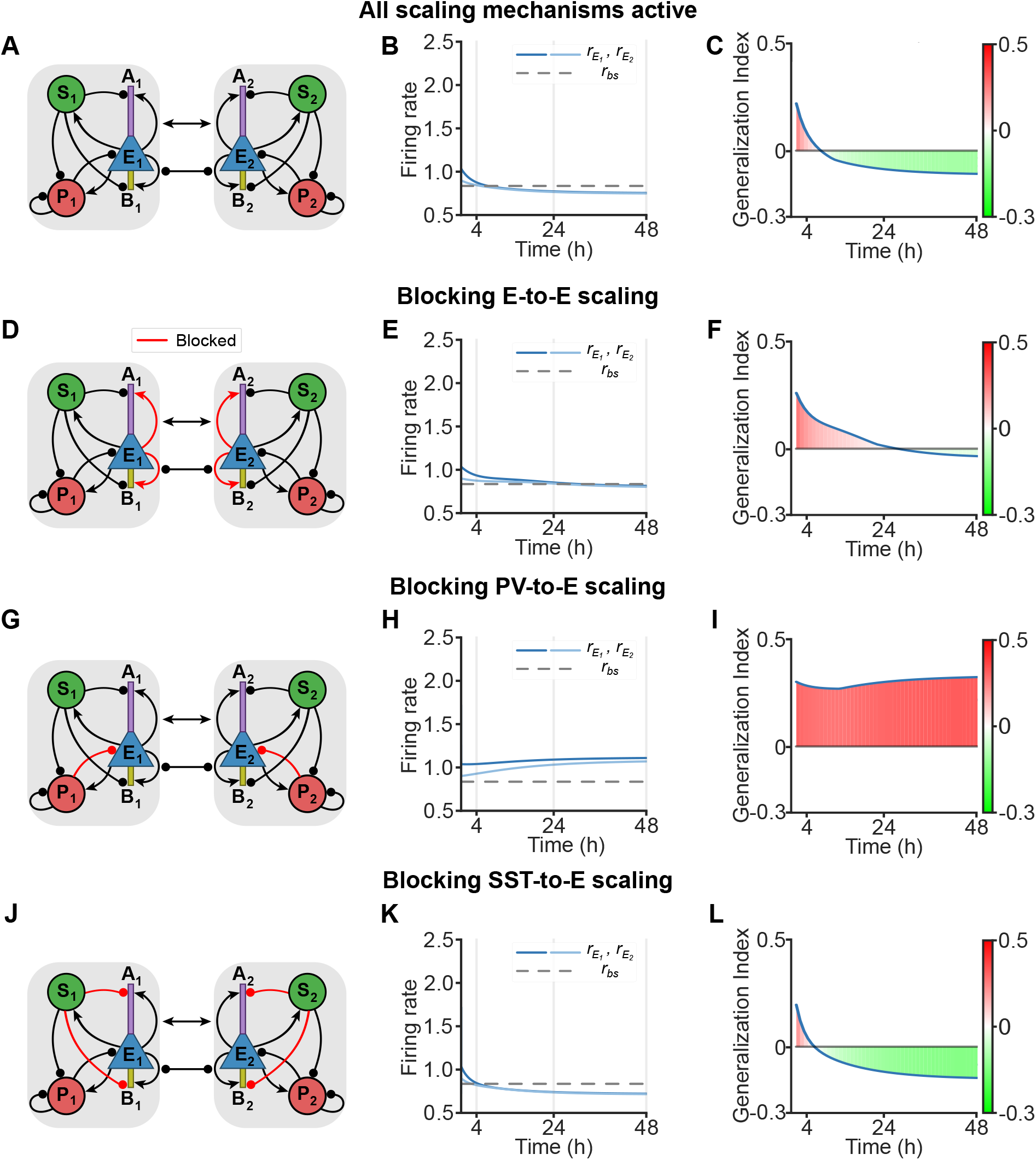
Multi-compartment model confirms cell-type-specific synaptic scaling to associative learning. **A**. Network schematic illustrating excitatory populations composed of three distinct compartments: soma (E, blue), apical den-drite (A, purple), and basal dendrite (B, yellow). PV (red) populations target the excitatory soma, whereas SST (green) populations target both apical and basal dendrites. **B**. Activity of excitatory population in subnetwork 1 (*r*_*E*1_) and sub-network 2 (*r*_*E*2_) after conditioning when blocking all scaling mechanisms. The gray horizontal dashed line represents the baseline activity level measured before conditioning. **C**. Evolution of the Generalization Index (GI) after conditioning when blocking all scaling mechanisms. A positive GI indicates memory generalization, whereas a negative GI represents memory specificity. **D - F**. Same as (A - C) but for blocking E-to-E scaling. **G - I**. Same as (A - C) but for blocking soma-targeting inhibitory scaling (e.g. PV-to-E scaling). **J - L**. Same as (A - C) but for blocking dendrite-targeting inhibitory scaling (e.g. SST-to-E scaling).

During conditioning, CS provides additional inputs to the basal dendrites of the excitatory populations and PV populations in subnetwork 1 (Figure S7). In the presence of the US, excitatory connections from the soma to both the apical and basal dendrites strengthen via Hebbian plasticity (Figure S7). Following conditioning, the network gradually transitions from memory generalization to memory specificity over time (Figure 8A-C), accompanied by downscaling of excitatory-to-excitatory connections, upscaling of soma-targeting inhibitory connections, and downscaling of dendrite-targeting inhibitory connections (Figure S7).

Consistent with the previous point neuron model, blocking excitatory-to-excitatory synaptic scaling delays the transition to memory specificity (Figure 8D-F), blocking soma-targeting inhibitory synaptic scaling prevents the formation of memory specificity (Figure 8G-I), and blocking dendritetargeting inhibitory synaptic scaling accelerates the achievement of memory specificity (Figure 8J-L). Taken together, this more biologically realistic model confirms the different contributions of cell-type-specific synaptic scaling mechanisms to associative learning.

## Discussion

Here, we investigated how different plasticity mechanisms shape associative learning in recurrent circuits comprising multiple interneuron types. Using analytical and computational approaches, we demonstrated that brief conditioning induces memory generalization through Hebbian plasticity. Following conditioning, different forms of synaptic scaling progressively establish memory specificity over time. Specifically, E-to-E and PV-to-E scaling function synergistically, but counteract SST-to-E scaling, to collectively govern the timing of memory refinement. Our findings reveal the cell-type-specific contributions of synaptic scaling and propose a role for top-down modulation in regulating associative learning.

Our study revealed several key insights into the mechanisms and consequences of associative learning. We demonstrated that different forms of synaptic scaling – a relatively slow process – are crucial for establishing memory specificity. This finding aligns with experimental observations showing that memory specificity emerges only several hours after conditioning (Wu et al., 2021). In the context of conditioned taste aversion, over time, the gradual fading of memory generalization may reduce food avoidance along with increasing hunger. Faster mechanisms, such as Hebbian inhibitory plasticity, could accelerate the elimination of food avoidance, but at the cost of a higher risk of encountering aversive food. In contrast, slower mechanisms, like synaptic scaling, may be more beneficial for animals to minimize risk while avoiding starvation. In addition to the gustatory cortex, similar associative learning paradigms have been applied in other sensory cortical regions (Letzkus et al., 2011; Pakan et al., 2018; Dalmay et al., 2019; Ottenheimer et al., 2023). Particularly, our model indicates that when all synaptic scaling mechanisms are intact, memory generalization occurs within 24 hours after conditioning (Figure 4B). Indeed, this is consistent with studies in the auditory cortex (Dalmay et al., 2019), which show that one or two days after conditioning, the evoked responses to CS-, corresponding to the test stimulus in our study, are lower than those observed during habituation without conditioning, which corresponds to the reference activity *r* ^*ref*^ in our model. Given the ubiquity of the cortical circuit motifs we modeled (Tremblay et al., 2016), our findings have the potential to provide broad insights into associative learning across sensory cortical regions.

Going beyond capturing existing experimental data, our model proposes a critical role of top-down influences in associative learning. Our findings demonstrate that a global, unspecific top-down signal, mediated by the unconditioned stimulus (e.g., punishment or reward), acts as a gate for the Hebbian learning process. In addition to these global signals, more specific top-down inputs, such as those related to attention and that target particular cell types (Park et al., 2025), can profoundly influence neural activity. These specific inputs can flexibly shift different subnetworks either into a long-term potentiation (LTP)-dominated or long-term depression (LTD)-dominated regime, thereby shaping associative learning and the timing of memory specificity emergence. Notably, while both types of inputs affect learning, global, unspecific top-down signals exert minimal influence on activity levels, whereas more specific, cell-type-targeted inputs affect learning by strongly modulating activity. Our model thus highlights that distinct sources of top-down inputs can act in parallel, each contributing to learning in mechanistically different ways.

Furthermore, our computational model allowed us to test and identify the cell-type-specific contributions to associative learning. Specifically, E-to-E scaling and PV-to-E scaling operate synergistically while opposing SST-to-E scaling. Our findings indicate that disabling excitatory synaptic scaling while preserving all forms of inhibitory synaptic scaling achieves memory specificity within 48 hours after conditioning. In contrast, disabling PV-to-E synaptic scaling while maintaining other scaling mechanisms prevents the establishment of memory specificity within the same period. These results highlight a powerful role of different forms of inhibitory synaptic scaling in facilitating precise associative learning. To preserve computational and analytical tractability, we made several simplifications. Biological neurons possess complex morphologies and exhibit a non-uniform distribution of ion channels and synaptic inputs across their dendritic trees (Jiang et al., 2015; Peng et al., 2021; Dorsett et al., 2021; Schneider-Mizell et al., 2025). These dendritic nonlinearities and localized synaptic interactions play a crucial role in integrating synaptic inputs and shaping neuronal output (Poirazi et al., 2003; London and Häusser, 2005; Larkum et al., 2009; Morabito et al., 2025), influencing network dynamics and learning processes. Furthermore, biological neurons communicate through discrete spikes, and the precise timing of these spikes can affect temporal coding and shape synaptic modifications via spike-timing-dependent plasticity (Bi and Poo, 1998; Song et al., 2000; Pfister and Gerstner, 2006). However, in the asynchronous irregular regime, characterized by irregular firing, weak pairwise correlations, and widely considered a typical operating state of the cortex, the macroscopic behavior of spiking networks is often well approximated by their firing rates (Vreeswijk and Sompolinsky, 1998). Therefore, although our model abstracts away several biological details, by using point neuron models combined with the incorporation of known connectivity properties and multi-compartment models with passive dendrites, our work provides valuable insight into how different synaptic scaling mechanisms influence associative learning.

In our work, we assume that the unconditioned stimulus (US) gates Hebbian learning without directly altering neural activity. Biologically, however, the US likely elicits an unconditioned response by activating certain neurons in sensory cortices. For example, in classical fear-conditioning, an electrical shock can modulate neural activity in the auditory cortex (Letzkus et al., 2011; Moczulska et al., 2013). It remains unclear whether US-active neurons in the sensory cortices can serve as a behavioral readout for probing memory generalization in the brain, or if this role is restricted to US-active neurons in downstream brain regions. In a computational model, however, US-active neurons can reasonably be used for this purpose, if explicitly modeled. In this case, generalized memory is indicated when the test stimulus evokes activity not only in neurons tuned to the test stimulus but also in some US-active neurons. In contrast, specific memory is indicated when the test stimulus evokes activity only in neurons tuned to the test stimulus, without engaging US-active neurons. Since US-active neurons are not explicitly modeled in our study, the activity evoked by the test stimulus is used directly as a proxy for the potential to recruit US-active neurons, and thus for assessing memory specificity. This approach enables us to evaluate memory specificity without introducing additional neural populations or parameters, thereby greatly reducing model complexity. Furthermore, adopting an alternative formulation of Hebbian plasticity in which both pre- and postsynaptic activities are baseline-subtracted does not alter our results, indicating that our findings are robust to different implementations of Hebbian plasticity (Figure S8).

Here, we primarily investigated various synaptic scaling mechanisms during associative learning while excluding long-term inhibitory Hebbian plasticity. Although inhibitory synapses are known to undergo modifications driven by Hebbian plasticity (Froemke et al., 2007; D’amour and Froemke, 2015; Hennequin et al., 2017; Lagzi et al., 2021; Schulz et al., 2021; Wu et al., 2022; Miehl and Gjorgjieva, 2022; Festa et al., 2025), experiments suggest that the timescale of long-term inhibitory plasticity might be too rapid to explain the prolonged duration of memory generalization and the gradual emergence of memory specificity at 48 hours after blocking E-to-E scaling (Wu et al., 2021). Therefore, we postulate that this delayed emergence of memory specificity is likely driven by slower processes, such as inhibitory synaptic scaling, as proposed in our study.

In addition, beyond the three cell types (E, PV, and SST) included in our model, several other inhibitory interneuron subtypes have been identified (Wilmes and Clopath, 2019; Hertäg and Sprekeler, 2020; Pardi et al., 2020; Canto-Bustos et al., 2022; Veit et al., 2023; Palmigiano et al., 2023; Hartung et al., 2024; Naumann et al., 2025). Among these, vasoactive intestinal peptide (VIP)-expressing interneurons are a prominent class often incorporated into canonical microcircuit motifs (Pfeffer et al., 2013; Waitzmann et al., 2024; Wu et al., 2026). VIP interneurons primarily inhibit SST cells and are known to receive top-down inputs, which can significantly impact recurrent network dynamics (Fu et al., 2014; Zhang et al., 2014; Dipoppa et al., 2018; Garrett et al., 2020; Bastos et al., 2023; Furutachi et al., 2024). Although VIP interneurons were not explicitly modeled in our study, by providing dedicated inputs to SST to emulate top-down modulation, our results suggest a pivotal role of top-down modulation in shaping associative learning. Furthermore, in addition to SST, top-down inputs can target other cell types within local recurrent circuits (Makino and Komiyama, 2015). By applying top-down inputs to both E and SST populations, we show that excitatory top-down input onto E neurons accelerates the emergence of memory specificity, whereas excitatory top-down input onto SST neurons delays it (Figure S9). This is consistent with the notion that top-down input to E neurons directly enhances E activity, whereas top-down input to SST neurons suppresses E activity through inhibitory pathways. Biologically, distinct cognitive and behavioral states, such as attention (Zhang et al., 2014), arousal (Niell and Stryker, 2010; Reimer et al., 2014; Vinck et al., 2015), and locomotion, (Fu et al., 2014; Erisken et al., 2014; Leinweber et al., 2017) are likely associated with different top-down pathways. These inputs may operate independently and preferentially target specific cell types. Future work will be needed to implement biologically informed connectivity for distinct top-down inputs and determine how they synergistically shape associative learning.

Together, our work offers new insights into how distinct plasticity mechanisms interact to shape associative learning, highlights the significant impact of top-down influences and synaptic scaling, and reveals the cell-type-specific contributions to the establishment of precise memory representations.

## Methods

### Rate-based population model

To investigate the role of cell-type-specific synaptic scaling in associative learning, we constructed a rate-based population model comprising two subnetworks. Each subnetwork includes one excitatory, one PV, and one SST population. Different subnetworks are tuned to different stimuli corresponding to different tastants in the conditioned taste aversion experiments. The dynamics of the network can be described as follows (Richter and Gjorgjieva, 2022):

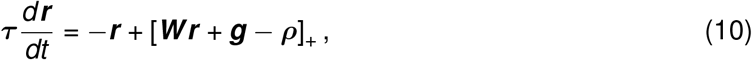

where ***τ*** is a diagonal matrix containing the time constants of firing rate dynamics for different populations, ***r*** is a vector containing the firing rates of different populations, ***g*** is a vector containing the inputs to different populations, and ***ρ*** is a vector containing the rheobases of different populations, []_+_ is a rectified function.

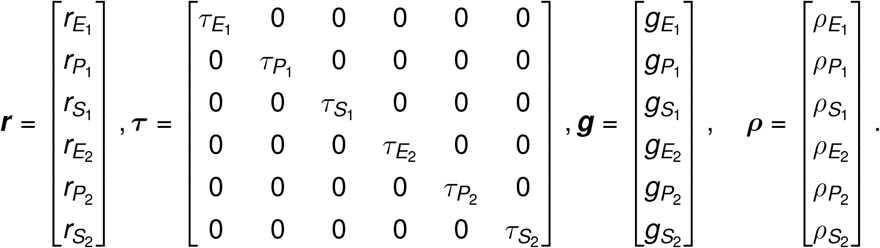

***W*** is the connectivity matrix defined as follows:

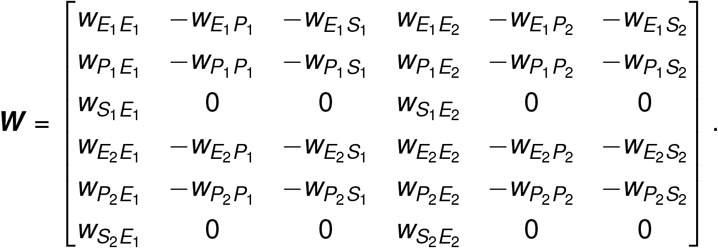

Before conditioning, the network settles into a steady state. To model the conditioned taste aversion experimental paradigm, specifically, to simulate the conditioned stimulus, additional inputs are provided to the excitatory (E_1_) and PV (P_1_) populations in the subnetwork 1 via increasing 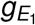 and 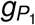 by *g*_Δ*E*_ and *g*_Δ*P*_, respectively. Similarly, to simulate the test stimulus, additional inputs are applied to the excitatory (E_2_) and PV (P_2_) populations in the subnetwork 2 via increasing 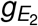 and 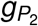 by *g*_Δ*E*_ and *g*_Δ*P*_, respectively. Parameter values for two subnetworks are the same unless mentioned otherwise.

### Three-factor Hebbian plasticity

Motivated by experimental studies showing that reward or punishment plays a decisive role in learning (Pawlak, 2010; Yagishita et al., 2014; He et al., 2015; Gerstner et al., 2018), we modeled Hebbian plasticity using a three-factor learning rule as follows:

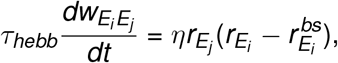

where *τ*_*hebb*_ is the time constant of Hebbian plasticity, 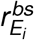 represents the baseline activity of the excitatory population in subnetwork *i* before conditioning, and *i, j* ∈ *{*1, 2*}*, representing the indices of subnetworks.

The third factor *η* is determined by the presence of the unconditioned aversive stimulus. More specifically,

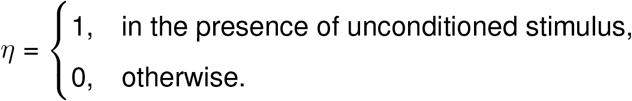

Thus, the third factor serves as a gate for Hebbian plasticity, enabling it during the conditioning phase while disabling it elsewhere.

### Synaptic scaling

The dynamics of the connection strength governed by synaptic scaling from the excitatory population in subnetwork *j* to the excitatory population in subnetwork *i* is given by (Van Rossum et al., 2000):

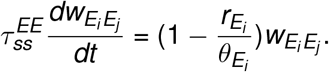

Similarly, for PV-to-E synaptic scaling, we have:

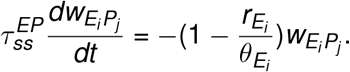

And for SST-to-E synaptic scaling, we have:

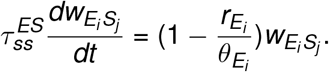

Here, *τ*_*ss*_ represents the time constant of synaptic scaling for individual type of connections, 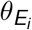 denotes the target firing rate or the set point of the excitatory population in the subnetwork *i*.

### Plasticity of set points

Set points of excitatory populations are subject to plastic changes, governed by the following dynamics:

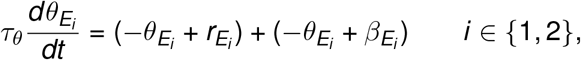

where *τ*_*θ*_ is the time constant governing the plasticity of the set points. The set point 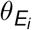 evolves based on the current activity 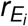 and the set point regulator 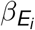 . The set point regulator *β* is dynamically updated according to:

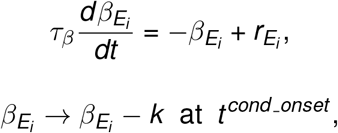

where *τ*_*β*_ denotes the time constant governing the plasticity of the set point regulator and *k* is a free parameter that determines the magnitude of the abrupt decrease in the set point regulator *β* of excitatory populations in both subnetworks at the onset of conditioning. Conditioning raises the activity *r*_*E*_, thereby increasing both the set point *θ* and the set point regulator *β*, in contrast, this sudden reduction in *β* counteracts the increases induced by conditioning and functions as a homeostatic mechanism to globally regulate the overall activity level (Kaleb et al., 2021).

### Analytical procedure

To thoroughly characterize the temporal evolution of memory specificity and generalization – specifically, how the network responds to the test stimulus following conditioning – we introduced a procedure to determine how set point regulators, set points, weights, and rates during conditioning and after conditioning evolve dynamically. In this procedure (Figure 4A), we defined two phases, and assumed that the firing rates of excitatory populations during conditioning (Phase 1) and after conditioning (Phase 2) in the absence of the test stimulus, which are experimentally measurable, are known. First, we formulate the time-variant firing rate of the excitatory population in the subnetwork *i* during conditioning (Phase 1) as an exponential function as follows:

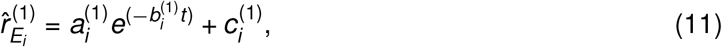

where 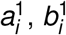, and 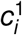 are coefficients obtained by fitting the parameterized functions to the excitatory firing rates of the subnetwork *i* during conditioning in the simulation. Superscripts ‘(1)’ indicate Phase 1 corresponding to the conditioning period.

By solving the three-factor Hebbian learning equation (Eq. 1), we can obtain 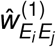 during conditioning as:

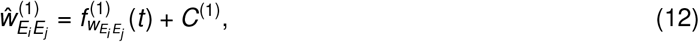

where

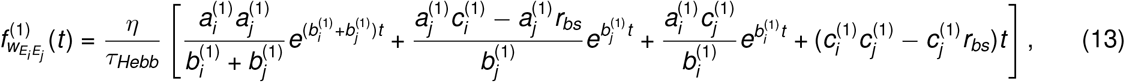

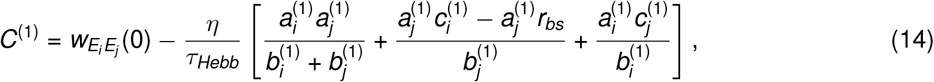

where 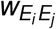 (0) is the initial value before conditioning. Note that here we disregarded the effects of synaptic scaling on the weights during conditioning due to its relatively slow timescale compared to the duration of the conditioning.

We then approximate the excitatory firing rates after conditioning (Phase 2) in a similar way:

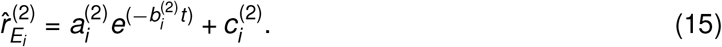

Given *k*, which represents the magnitude of the abrupt decrease in the set point regulator *β*, the set point regulator 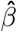 after conditioning can be determined analytically as follows:

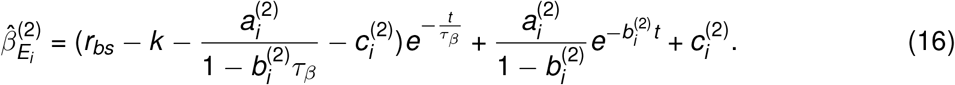

Using the 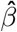 from the above equation, the set points of excitatory populations 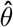 after conditioning can be calculated as follows:

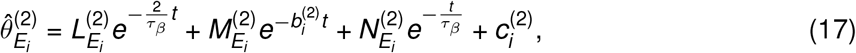

with

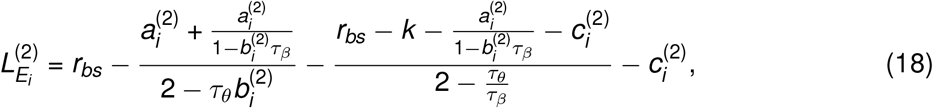

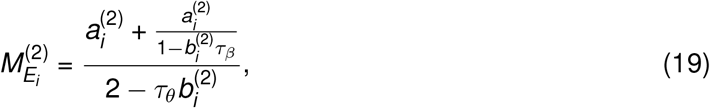

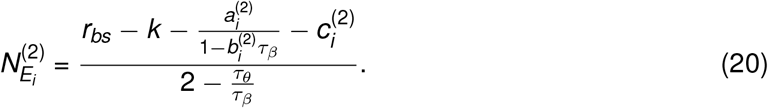

Using the above obtained 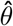, the weights after conditioning can be computed as follows:

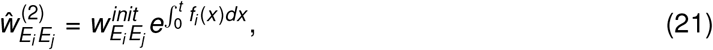

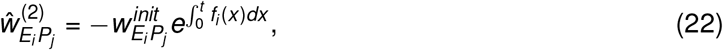

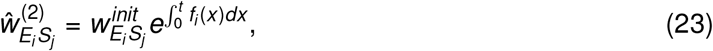

with

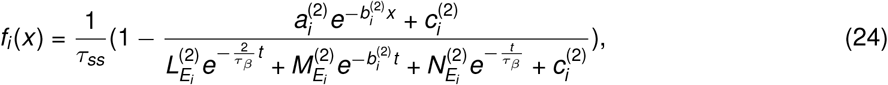

where 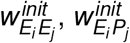, and 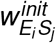 denote the weights right after conditioning from the excitatory, PV and SST population in the subnetwork *j* to the excitatory population in the subnetwork *i*. We obtain 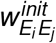 from Eq. 12. In contrast, 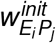 and 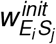 are identical to their initial values 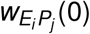 and 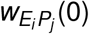, respectively, as they remain unchanged during conditioning.

From the above-obtained weights, assuming all populations exhibit non-zero firing rates, the steadystate firing rate for any given input at any given time after conditioning can be calculated as follows:

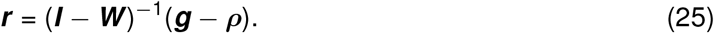

By substituting the inputs from the test stimulus condition into the above equation, we can determine the responses to the test stimulus over time, providing a comprehensive description of the evolution of memory specificity and generalization.

### Alternative form of three-factor Hebbian plasticity

To assess the robustness of our results, we implemented an alternative three-factor Hebbian learning rule in which baseline activity is subtracted from both the pre- and postsynaptic activity.

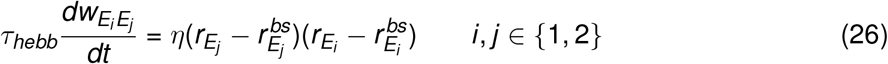

### Sensitivity analysis

To assess the robustness of our results to parameter selection, we conducted grid simulations across the model’s parameter space by changing the connection strengths to excitatory populations (E-to-E, PV-to-E, and SST-to-E, (Figure S4) and to inhibitory populations (E-to-PV, PV-to-PV, SST-to-PV, and E-to-SST, (Figure S5) both within the subnetwork and cross the subnetworks from 0.01 to 1.01 with increments of 0.1. To ensure computational feasibility, we further constrained the parameters for the cross-connections to be smaller than those for within connections (e.g. 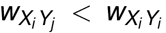), resulting in a total of 72728 simulations. We then performed these simulations and evaluated memory specificity in models with varying initial conditions of either connections to excitatory or inhibitory populations. We selected models that meet two experimental conditions: (1) a transition from memory generalization to memory specificity occurs between 4h and 24h, and (2) when E-to-E scaling is blocked, this transition occurs later than in (1). Models that meet these criteria are labeled as respecting ‘experimentally-matched models’ (in total 828 out of 58087 in Figure S4, 258 out of 14641 in Figure S5). Using these experimentally-matched models, we evaluated whether the role of cell-specific synaptic scaling mechanisms aligns with the instance presented in the Results Section (Figure 7) by blocking either PV-to-E or SST-to-E scaling. If blocking PV-to-E scaling diminishes memory specificity and blocking SST-to-E scaling accelerates the transition from memory generalization to memory specificity in these selected models, we labeled them as ‘aligned models’ (Figure S4, S5). We found that 96% of the selected models (1041 out of 1086) exhibit the same cell-type-specific contributions as in the presented example (Figure 7), along with a wide range of baseline firing rates (Figure S6), suggesting that our results are robust to parameter selection.

### Multi-compartment model

To verify that the conclusions drawn from the point neuron model remain valid in a more biologically detailed setting, we constructed a multi-compartment model comprising two subnetworks. In contrast to the earlier point neuron implementation, the excitatory population in this model consists of three distinct compartments: soma (E), apical dendrite (A), and basal dendrite (B). PV and SST populations are still modeled as point neurons, but they now target the soma and dendrites (both apical and basal dendrites) of the excitatory populations, respectively. As in the point neuron model, the two subnetworks are tuned to different stimuli, corresponding to distinct tastants in the conditioned taste aversion experiments. The firing rate dynamics of the excitatory populations are modeled as

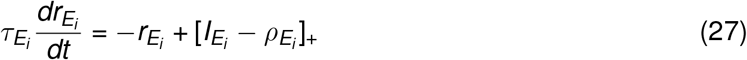

where 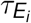 is the time constant of the dynamics, 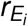 is the firing rate of the excitatory population in the subnetwork *i*, 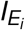 denotes the somatic current of the excitatory population in subnetwork *i*, and 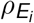 is its rheobase.

The somatic current of the excitatory population is given by:

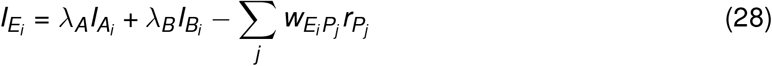

comprising contributions from apical dendritic currents 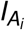 scaled by the coupling factor *λ*_*A*_, basal dendritic currents 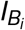 scaled by the coupling factor *λ*_*B*_, and soma-targeting inhibition from PV populations across both subnetworks.

The apical dendritic current is given by:

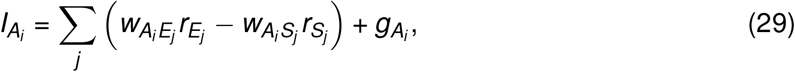

comprising recurrent excitatory inputs to the apical dendrites, SST-mediated dendritic inhibition, and apical dendrite-specific background input *g*_*A*_.

The basal dendritic current is given by:

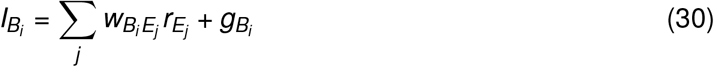

comprising recurrent excitatory inputs to the basal dendrites and basal dendrite-specific background input *g*_*B*_.

The firing rate dynamics of the PV populations are modeled as:

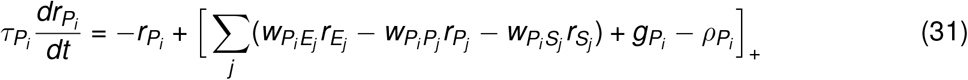

The firing rate dynamics of the SST populations are modeled as:

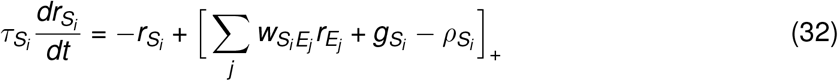

To model the conditioned taste aversion experimental paradigm, additional inputs are now provided to the basal dendrite of the excitatory population (*B*_1_) and the PV (*P*_1_) population in the subnetwork 1 via increasing 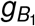 and 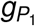 by 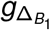 and 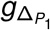, respectively. Similarly, to simulate the test stimulus, additional inputs are applied to the basal dendrite of the excitatory population (*B*_2_) and the PV (*P*_2_) population in the subnetwork 2 via increasing 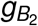 and 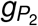 by 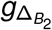 and 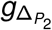, respectively.

During conditioning, the recurrent excitatory connections to the apical and basal dendrites of excitatory populations undergo Hebbian plasticity, governed by a three-factor Hebbian learning rule:

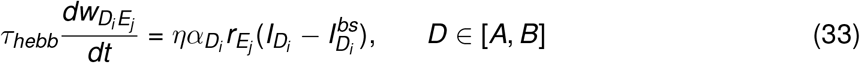

where *τ*_*hebb*_ is the time constant of Hebbian plasticity, *D* denotes either apical dendrite (*A*) or basal dendrite (*B*), 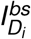 is the baseline current at the specified dendritic location in subnetwork *i* before conditioning, and *i, j* ∈ *{*1, 2*}* are the indices of subnetworks. The third factor, *η*, which serves as a gating signal for Hebbian plasticity, is defined as in the previous model. And 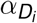 is a location-specific learning rate that determines the change in amplitude induced by Hebbian learning.

The dynamics of recurrent excitatory connections to the apical and basal dendrites of excitatory populations, governed by synaptic scaling, are given by:

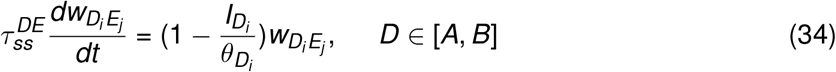

Similarly, for PV-mediated, soma-targeting inhibitory synaptic scaling, we have:

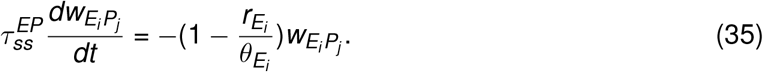

And for SST-mediated, dendrite-targeting inhibitory synaptic scaling, we have:

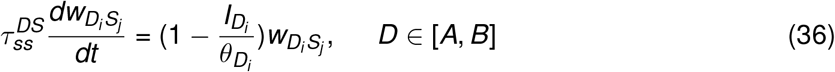

Here, *τ*_*ss*_ represents the time constant of synaptic scaling for individual type of connections, *θ* denotes the target activity level or the set point of the excitatory population 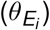, the apical dendrite 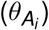, the basal dendrite 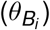 in the subnetwork *i*.

The set points, *θ*, and set point regulators, *β*, are defined as follows, in a manner similar to before (Eq.6-8):

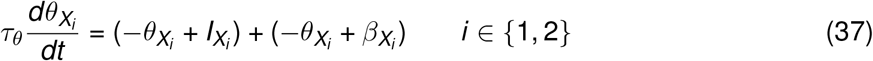

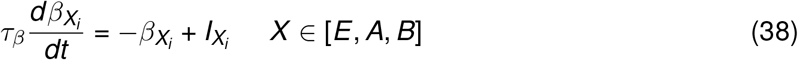

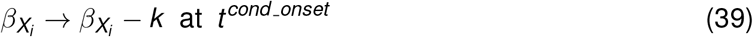

### Numerical simulation

Simulations were performed in Python with Numba (Lam et al., 2015). Differential equations were implemented by Euler method with a timestep of 0.1 milliseconds. All simulation parameters are listed in Table 1,2.

**Table 1.** Model Parameters.

| Network dynamics and network connectivity |  |  |  |
| --- | --- | --- | --- |
| Symbol | Value | Unit | Description |
| $\tau_E$ | 20 | ms | time constant of E rate dynamics |
| $\tau_P$ | 5 | ms | time constant of PV rate dynamics |
| $\tau_S$ | 10 | ms | time constant of SST rate dynamics |
| $W_{EE_{ii}}$ | 0.51 | a.u. | connection strength from E to E within subnetworks |
| $W_{EE_{ij}}$ | 0.51 | a.u. | connection strength from E to E across subnetworks |
| $W_{EP_{ii}}$ | 0.91 | a.u. | connection strength from PV to E within subnetworks |
| $W_{EP_{ij}}$ | 0.41 | a.u. | connection strength from PV to E across subnetworks |
| $W_{ES_{ii}}$ | 0.51 | a.u. | connection strength from SST to E within subnetworks |
| $W_{ES_{ij}}$ | 0.31 | a.u. | connection strength from SST to E across subnetworks |
| $W_{PE_{ii}}$ | 0.3 | a.u. | connection strength from E to PV within subnetworks |
| $W_{PE_{ij}}$ | 0.1 | a.u. | connection strength from E to PV across subnetworks |
| $W_{PP_{ii}}$ | 0.2 | a.u. | connection strength from PV to PV within subnetworks |
| $W_{PP_{ij}}$ | 0.1 | a.u. | connection strength from PV to PV across subnetworks |
| $W_{PS_{ii}}$ | 0.3 | a.u. | connection strength from SST to PV within subnetworks |
| $W_{PS_{ij}}$ | 0.1 | a.u. | connection strength from SST to PV across subnetworks |
| $W_{SE_{ii}}$ | 0.4 | a.u. | connection strength from E to SST within subnetworks |
| $W_{SE_{ij}}$ | 0.1 | a.u. | connection strength from E to SST across subnetworks |
| $\rho$ | 1.5 | a.u. | rheobase current for E, PV, and SST |
| Plasticity mechanisms |  |  |  |
| $\tau_{hebb}$ | 4 | min | time constant of three-factor Hebbian learning |
| $\tau_\theta$ | 24 | h | time constant of target activity dynamics |
| $\tau_\beta$ | 28 | h | time constant of target activity regulator dynamics |
| $\tau_{ss}$ | 9 | h | time constant of synaptic scaling dynamics (except Figure S8) |
| $\tau_{ss}$ | 11 | h | time constant of synaptic scaling dynamics (Figure S8) |
| $k$ | 0.25 | a.u. | amplitude of the decrease in target activity regulator |
| Inputs |  |  |  |
| $g_E$ | 4.5 | a.u. | background input to E |
| $g_P$ | 3.2 | a.u. | background input to PV |
| $g_S$ | 3 | a.u. | background input to SST |
| $g_{\Delta E}$ | 1 | a.u. | additional input to $E$ |
| $g_{\Delta P}$ | 0.5 | a.u. | additional input to $P$ |
| $g_{\Delta S}$ | 0.6 (−0.3) | a.u. | additional top-down excitatory (inhibitory) input to SST |

**Table 2.**
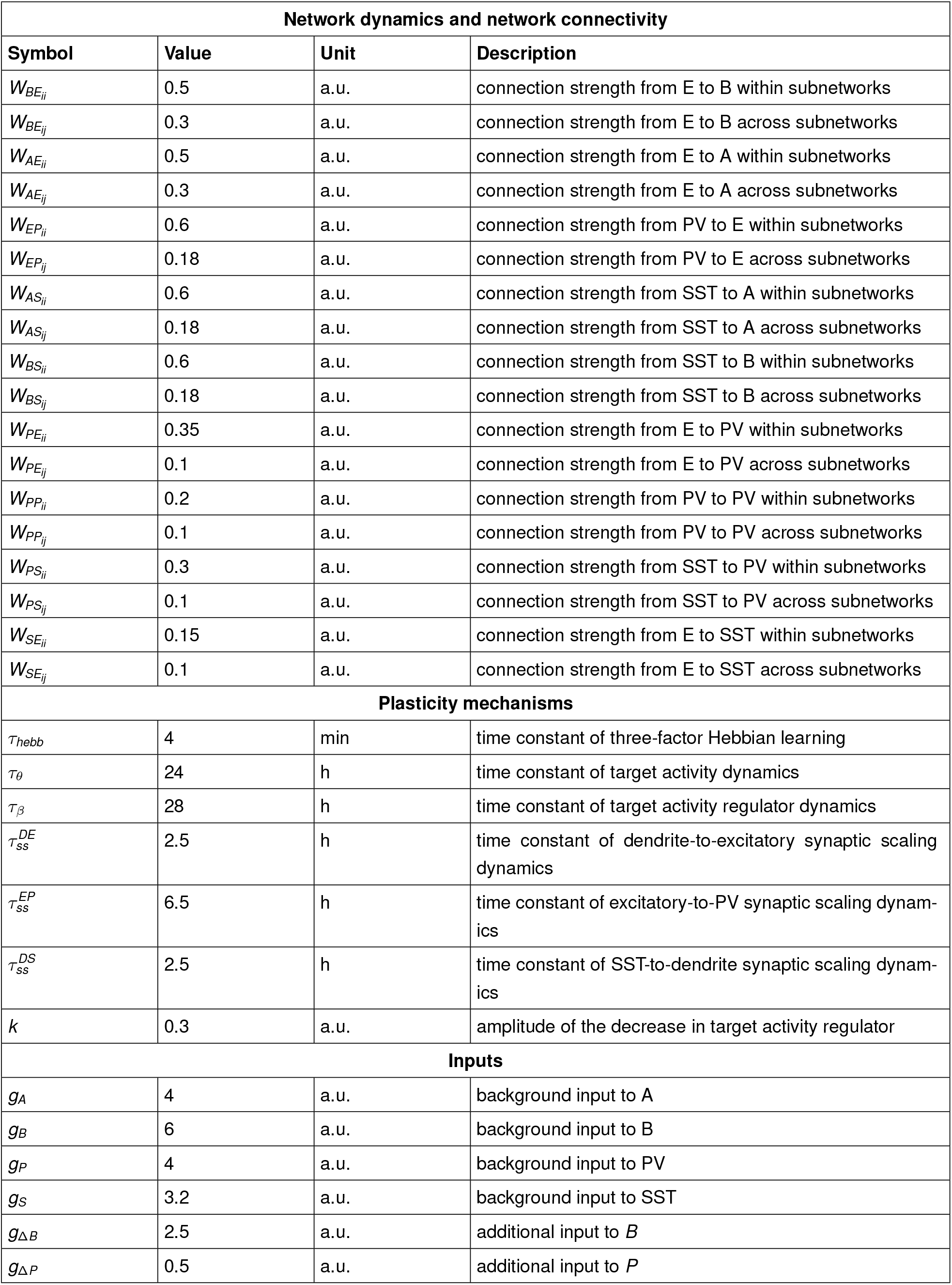
Parameters for multi-compartment model. *τ*_*E*_, *τ*_*P*_, *τ*_*S*_ and *ρ* are consistent with Table 1.

| Network dynamics and network connectivity |  |  |  |
| --- | --- | --- | --- |
| Symbol | Value | Unit | Description |
| $W_{BE_{ii}}$ | 0.5 | a.u. | connection strength from E to B within subnetworks |
| $W_{BE_{ij}}$ | 0.3 | a.u. | connection strength from E to B across subnetworks |
| $W_{AE_{ii}}$ | 0.5 | a.u. | connection strength from E to A within subnetworks |
| $W_{AE_{ij}}$ | 0.3 | a.u. | connection strength from E to A across subnetworks |
| $W_{EP_{ii}}$ | 0.6 | a.u. | connection strength from PV to E within subnetworks |
| $W_{EP_{ij}}$ | 0.18 | a.u. | connection strength from PV to E across subnetworks |
| $W_{AS_{ii}}$ | 0.6 | a.u. | connection strength from SST to A within subnetworks |
| $W_{AS_{ij}}$ | 0.18 | a.u. | connection strength from SST to A across subnetworks |
| $W_{BS_{ii}}$ | 0.6 | a.u. | connection strength from SST to B within subnetworks |
| $W_{BS_{ij}}$ | 0.18 | a.u. | connection strength from SST to B across subnetworks |
| $W_{PE_{ii}}$ | 0.35 | a.u. | connection strength from E to PV within subnetworks |
| $W_{PE_{ij}}$ | 0.1 | a.u. | connection strength from E to PV across subnetworks |
| $W_{PP_{ii}}$ | 0.2 | a.u. | connection strength from PV to PV within subnetworks |
| $W_{PP_{ij}}$ | 0.1 | a.u. | connection strength from PV to PV across subnetworks |
| $W_{PS_{ii}}$ | 0.3 | a.u. | connection strength from SST to PV within subnetworks |
| $W_{PS_{ij}}$ | 0.1 | a.u. | connection strength from SST to PV across subnetworks |
| $W_{SE_{ii}}$ | 0.15 | a.u. | connection strength from E to SST within subnetworks |
| $W_{SE_{ij}}$ | 0.1 | a.u. | connection strength from E to SST across subnetworks |
| Plasticity mechanisms |  |  |  |
| $\tau_{hebb}$ | 4 | min | time constant of three-factor Hebbian learning |
| $\tau_\theta$ | 24 | h | time constant of target activity dynamics |
| $\tau_\beta$ | 28 | h | time constant of target activity regulator dynamics |
| $\tau_{ss}^{DE}$ | 2.5 | h | time constant of dendrite-to-excitatory synaptic scaling dynamics |
| $\tau_{ss}^{EP}$ | 6.5 | h | time constant of excitatory-to-PV synaptic scaling dynamics |
| $\tau_{ss}^{DS}$ | 2.5 | h | time constant of SST-to-dendrite synaptic scaling dynamics |
| $k$ | 0.3 | a.u. | amplitude of the decrease in target activity regulator |
| Inputs |  |  |  |
| $g_A$ | 4 | a.u. | background input to A |
| $g_B$ | 6 | a.u. | background input to B |
| $g_P$ | 4 | a.u. | background input to PV |
| $g_S$ | 3.2 | a.u. | background input to SST |
| $g_{\Delta B}$ | 2.5 | a.u. | additional input to B |
| $g_{\Delta P}$ | 0.5 | a.u. | additional input to P |

## Data Availability

The simulation code is publicly available at https://github.com/comp-neural-circuits/cell-type-specific-synaptic-scaling.

## Contributions

Y.K.W. and J.G. designed research; F.V. and A.K. performed the numerical simulations; F.V. and Y.K.W. performed the analytic calculations; F.V., A.K., Y.K.W., and J.G. wrote the paper.

## Acknowledgments

We thank Gina Turrigiano and Chi-Hong Wu as well as members of the Computation in Neural Circuits Group for helpful discussions. This work was funded by the European Research Council (Grant Agreement No. 804824 to J.G.) and by the DFG in the Collaborative Research Centre 1080 (to J.G.). A.K. was also supported by TUM and the Elite Network of Bavaria. Y.K.W. was also supported by the Add-on Fellowship of the Joachim Herz Foundation.

## Supplementary Material

**Fig. S1.**
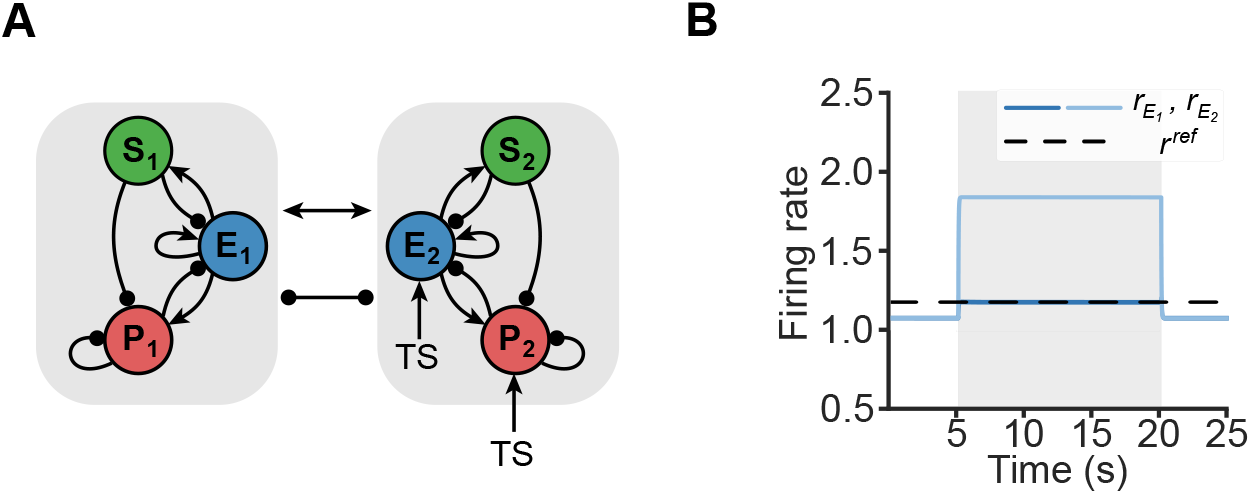
Definition of reference activity. **A**. Network schematic when applying a conditioned stimulus to subnetwork 2 in the absence of unconditioned stimulus. **B**. The reference activity *r* ^*ref*^ is determined by measuring the excitatory population activity of subnetwork 1 in response to the test stimulus in the absence of plasticity. The test stimulus is applied during the period marked in gray, and modeled by providing additional inputs to the excitatory and PV population of subnetwork 2.

**Fig. S2.**
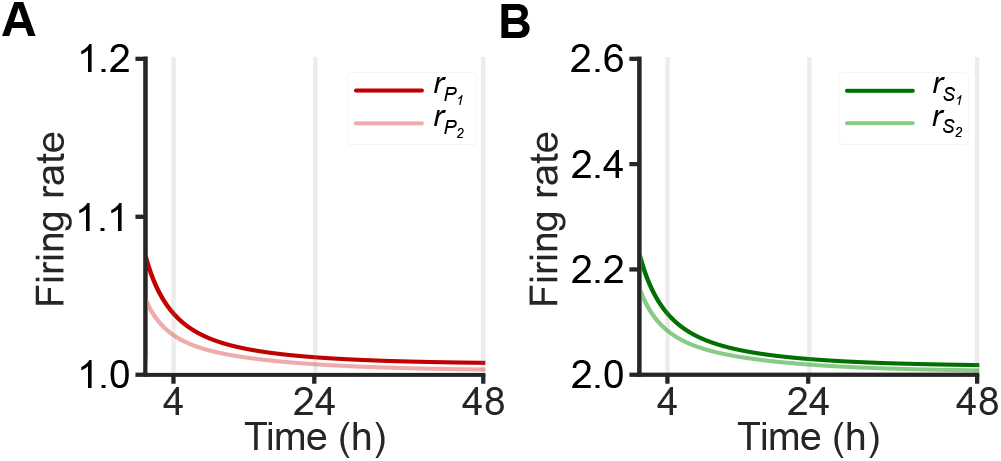
Inhibitory population activity post-conditioning. **A**. Activity of PV population in subnetwork 1 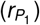 and subnet-work 2 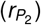 after conditioning. **B**. Same as A but for SST population.

**Fig. S3.**
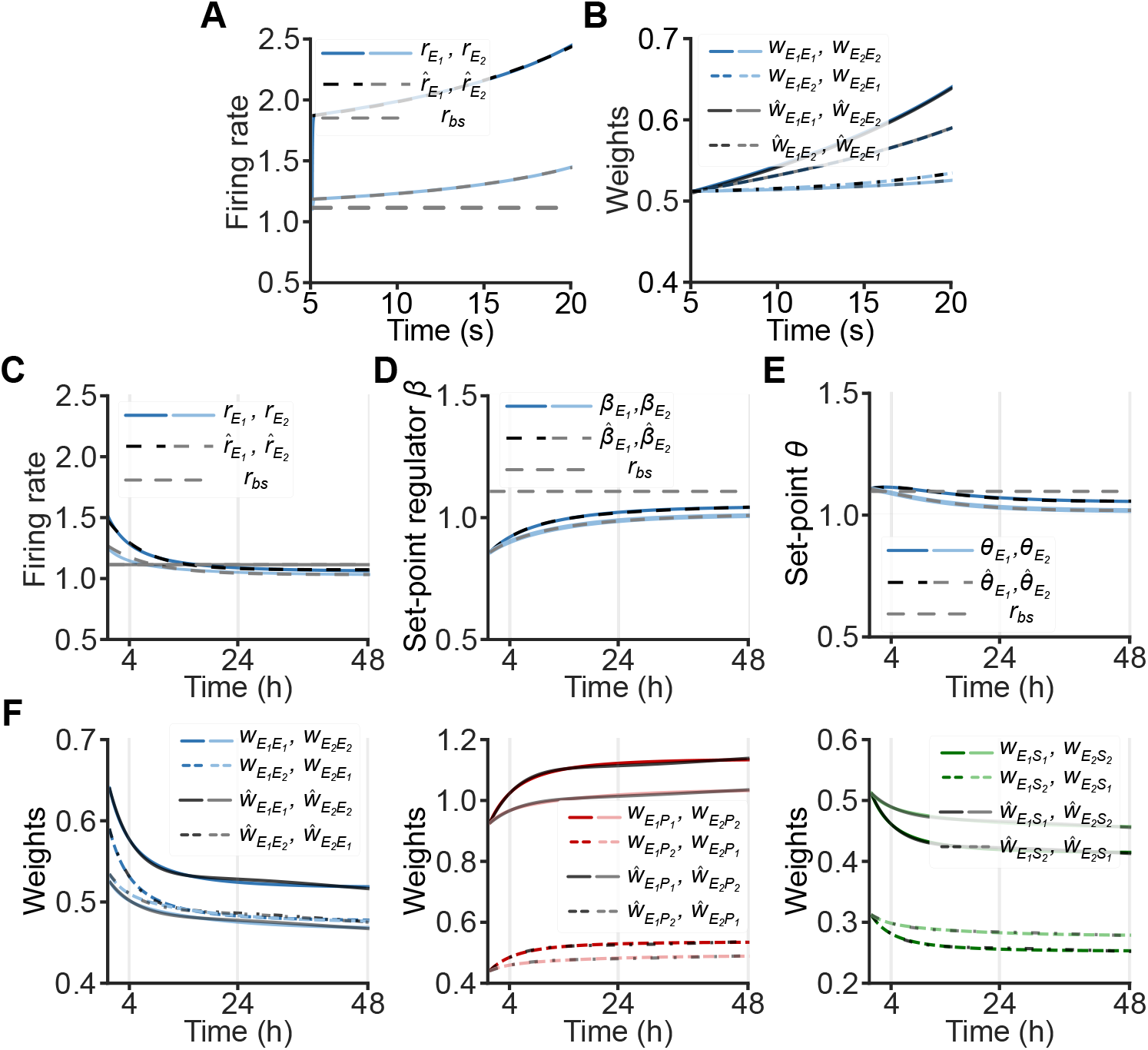
Comparisons between numerical simulations and analytical calculations during conditioning in Phase 1 and post-conditioning in Phase 2. **A**. Simulated excitatory activity of subnetwork 1 (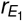, same as in Figure 2) and subnetwork 2 (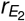, same as in Figure 2) during conditioning, and fitted excitatory activity of subnetwork 1 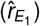 and subnetwork 2 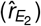 during conditioning. The horizontal dashed line represents the baseline activity level measured before conditioning. **B**. Numerical simulated excitatory weight evolution (same as in Figure 2) during conditioning, and analytical calculated excitatory weights evolution during conditioning. **C**. Simulated excitatory activity of subnetwork 1 (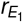, same as in Figure 3) and subnetwork 2 (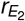, same as in Figure 3) after conditioning, and fitted excitatory activity of subnetwork 1 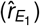 and subnetwork 2 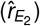 after conditioning. The horizontal dashed line represents the baseline activity level measured before conditioning. **D**. Same as C but for set point regulators after conditioning. **E**. Same as D but for set points after conditioning. **F**. (Left) Numerical simulated excitatory weight evolution (same as in Figure 3) after conditioning, and analytical calculated excitatory weights evolution after conditioning. (Middle, Right) Same as Left but for PV-to-E and SST-to-E weights respectively.

**Fig. S4.**
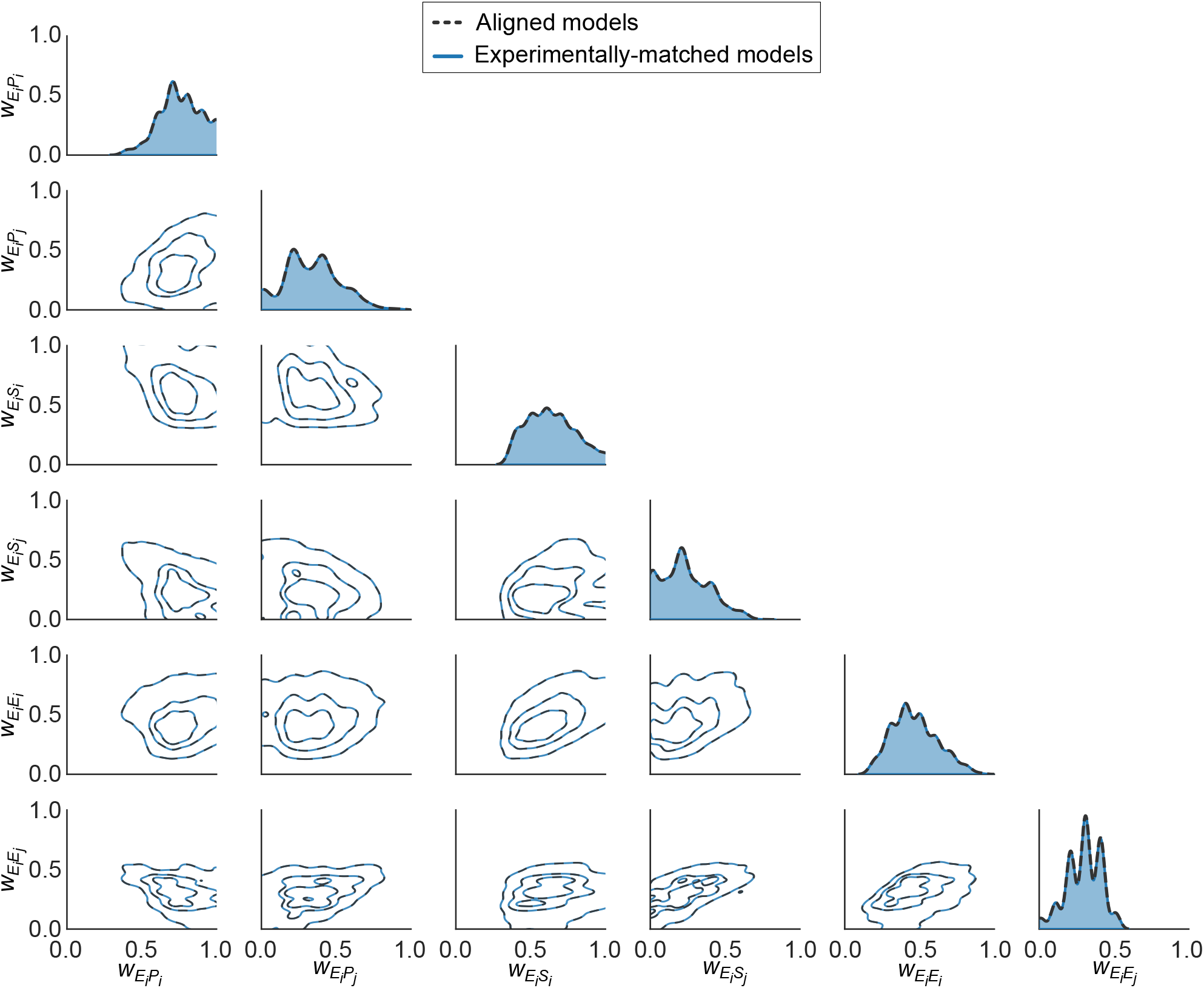
Pairwise density plots of plastic weight parameter distributions for experimentally matched models and aligned models. Experimentally matched models are defined as those in which the transition from memory generalization to specificity occurs between 4h and 24h when all scaling mechanisms are active and shifts later when E-to-E scaling is blocked. Aligned models are a subset of experimentally matched models, characterized by a diminished memory specificity when PV-to-E scaling is blocked and an accelerated transition when SST-to-E scaling is blocked. The x- and y-axes represent specific plastic connection weights. The substantial overlap between the two distributions suggests that cell-type-specific contributions to associative learning demonstrated in Figure 7 are robust to parameter selection. Here, 98% of the experimentally matched models (811 out of 828) are aligned models, hence many of the contour lines lie on top of each other.

**Fig. S5.**
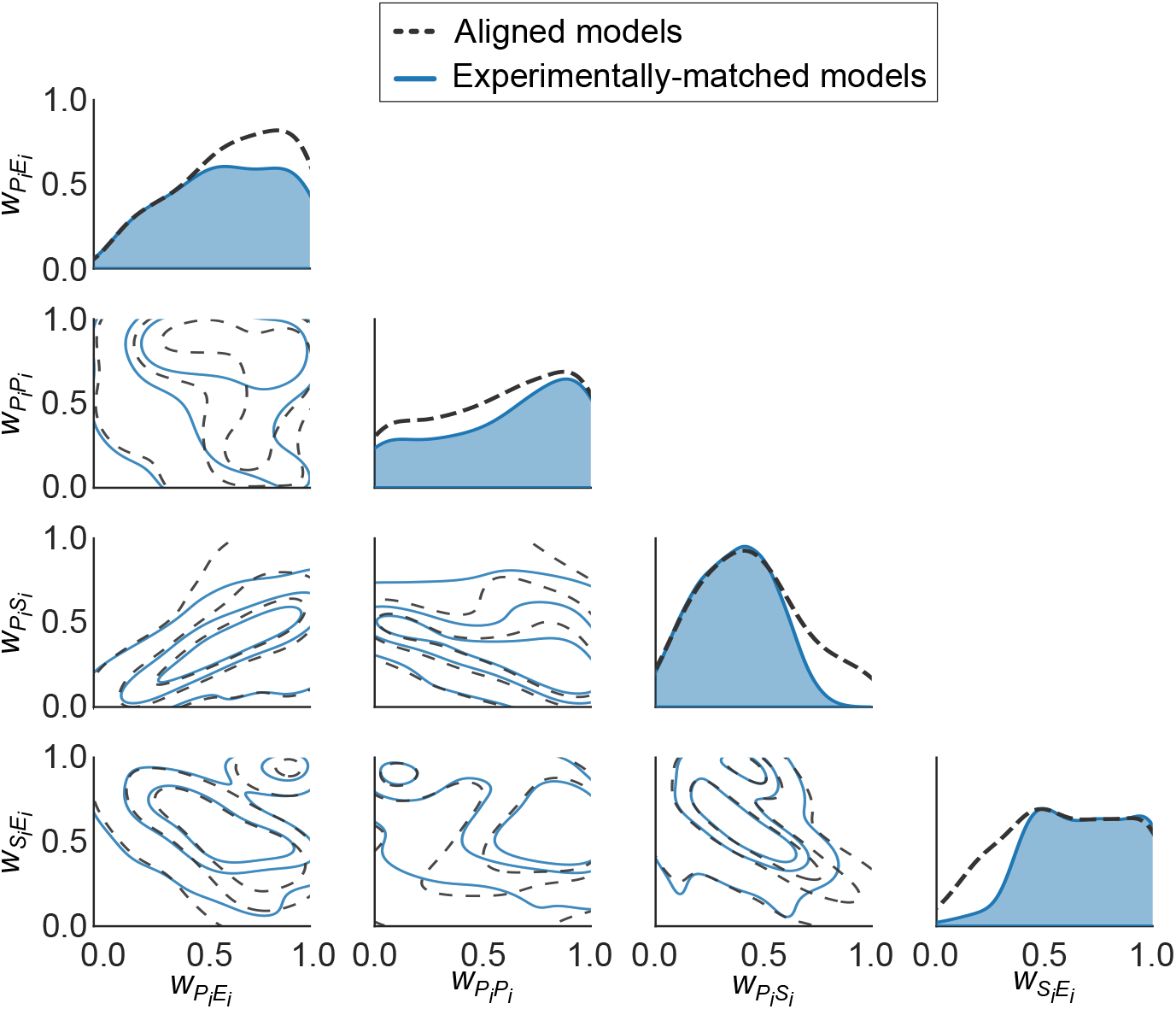
Pairwise density plots of static weight parameter distributions for experimentally matched models and aligned models. Experimentally matched models are defined as those in which the transition from memory generalization to specificity occurs between 4h and 24h when all scaling mechanisms are active and shifts later when E-to-E scaling is blocked. Aligned models are a subset of experimentally matched models, characterized by a diminished memory specificity when PV-to-E scaling is blocked and an accelerated transition when SST-to-E scaling is blocked. The x- and y-axes represent specific static connection weights. The substantial overlap between the two distributions suggests that cell-type-specific contributions to associative learning demonstrated in Figure 7 are robust to parameter selection. Here, 89% of the experimentally matched models (230 out of 258) are aligned models.

**Fig. S6.**
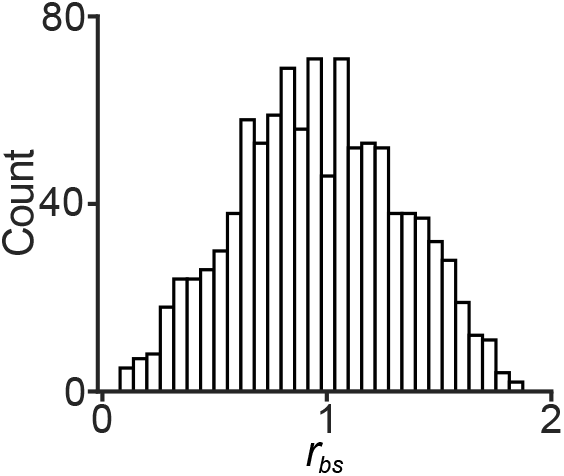
Baseline activity (*r*_*bs*_) histogram of all aligned models in Fig. S4 and S5. The broad distribution suggests that the exact value of the baseline activity is not important to capture the model’s results.

**Fig. S7.**
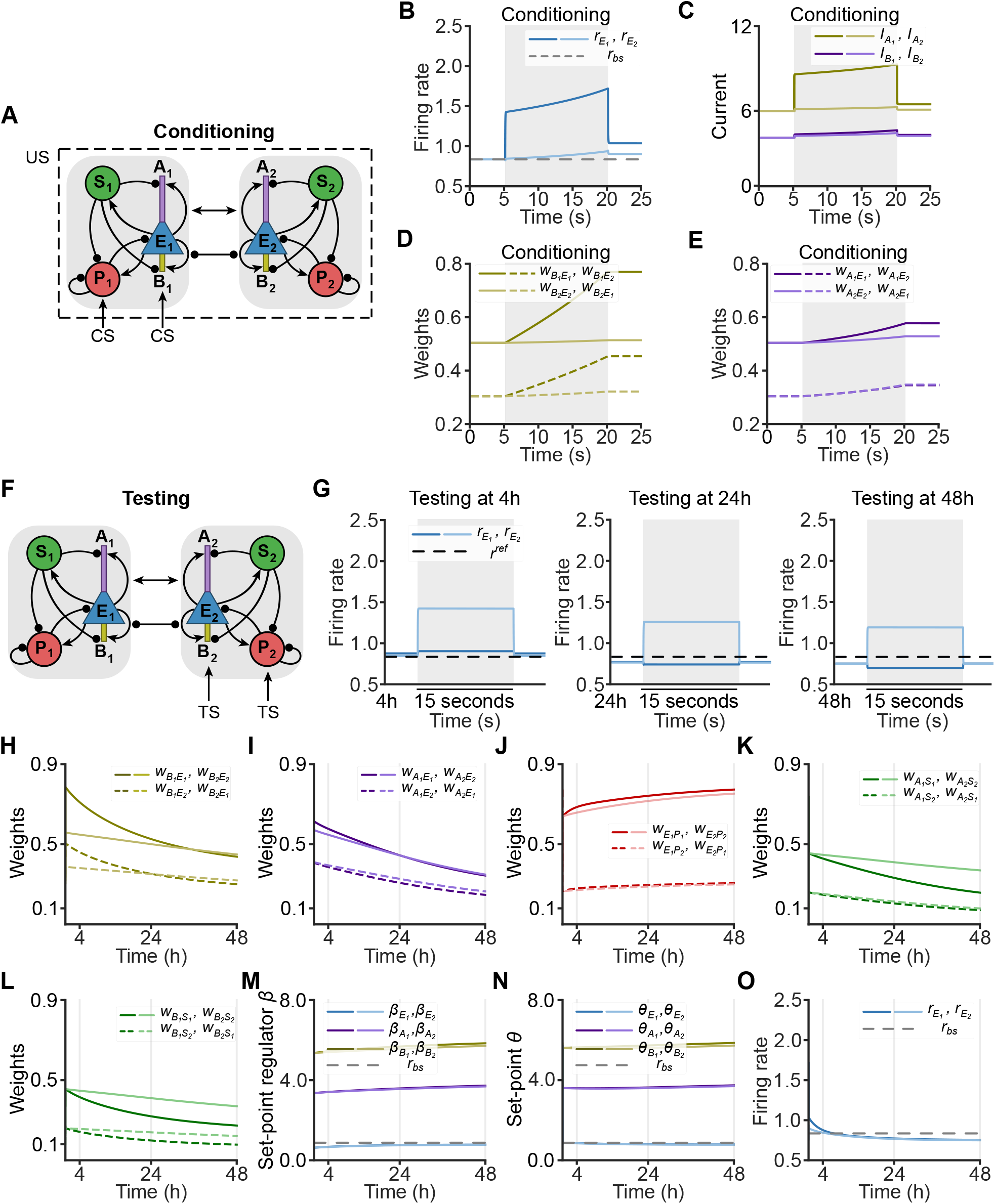
Multi-compartment model during and after conditioning. **A**. Network schematic of the conditioning phase. In the multi-compartment model, excitatory populations are composed of three distinct compartments: soma (E, blue), apical dendrite (A, purple), and basal dendrite (B, yellow). PV (red) populations target the excitatory soma, whereas SST (green) populations target both apical and basal dendrites. During conditioning, the basal dendrite of the excitatory population and PV population of subnetwork 1 receive additional inputs that correspond to the conditioned stimulus. During this phase, the unconditioned stimulus is present, modulating Hebbian plasticity. **B**. Activity of excitatory population in subnetwork 1 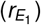 and subnetwork 2 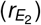 during conditioning. Conditioning is applied during the interval from 5 to 20s by increasing the inputs to the basal dendrite (B1) and the PV (P1) populations in subnetwork 1. The dashed line represents the baseline activity level measured before conditioning. **C**. Apical dendritic currents (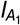 and 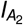) and basal dendritic currents (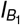and 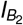) in subnetwork 1 and 2. Different types of currents are indicated by the differently colored lines. **D**. Excitatory-to-basal dendrite connection strength during conditioning. Different connections are indicated by the differently colored lines. **E**. Same as D but for excitatory-to-apical dendrite connection strength. **F**. Network schematic of the testing phase. After conditioning, the basal dendrite (B2) and the PV (P2) population of subnetwork 2 receive additional inputs that correspond to the test stimulus. **G**. Responses of excitatory populations in subnetwork 1 and subnetwork 2 when presenting a test stimulus for 15s (gray) at 4h (left), 24h (middle) and 48h (right). The black horizontal lines indicate the reference activity (*r* ^*ref*^), measured by the excitatory population in subnetwork 1 in response to a test stimulus under identical initial weights conditions but without plasticity. **H**. Excitatory-to-basal dendrite connection strength after conditioning. Different connections are indicated by the differently colored lines. **I**. Same as H but for excitatory-to-apical dendrite connection strength. **J**. Same as H but for PV-to-excitatory soma connection strength. **K**. Same as H but for SST-to-excitatory apical dendrite connection strength. **L**. Same as H but for SST-to-excitatory basal dendrite connection strength. **M**. Evolution of set point regulators for excitatory soma activity (E, blue), apical dendritic activity (A, purple), and basal dendritic activity (B, yellow) in subnetworks 1 and 2 after conditioning up to 48h. The gray horizontal dashed line represents the corresponding baseline activity level measured before conditioning. **N**. Same as L but for the corresponding set points *θ*. **O**. Activity of excitatory populations after conditioning.

**Fig. S8.**
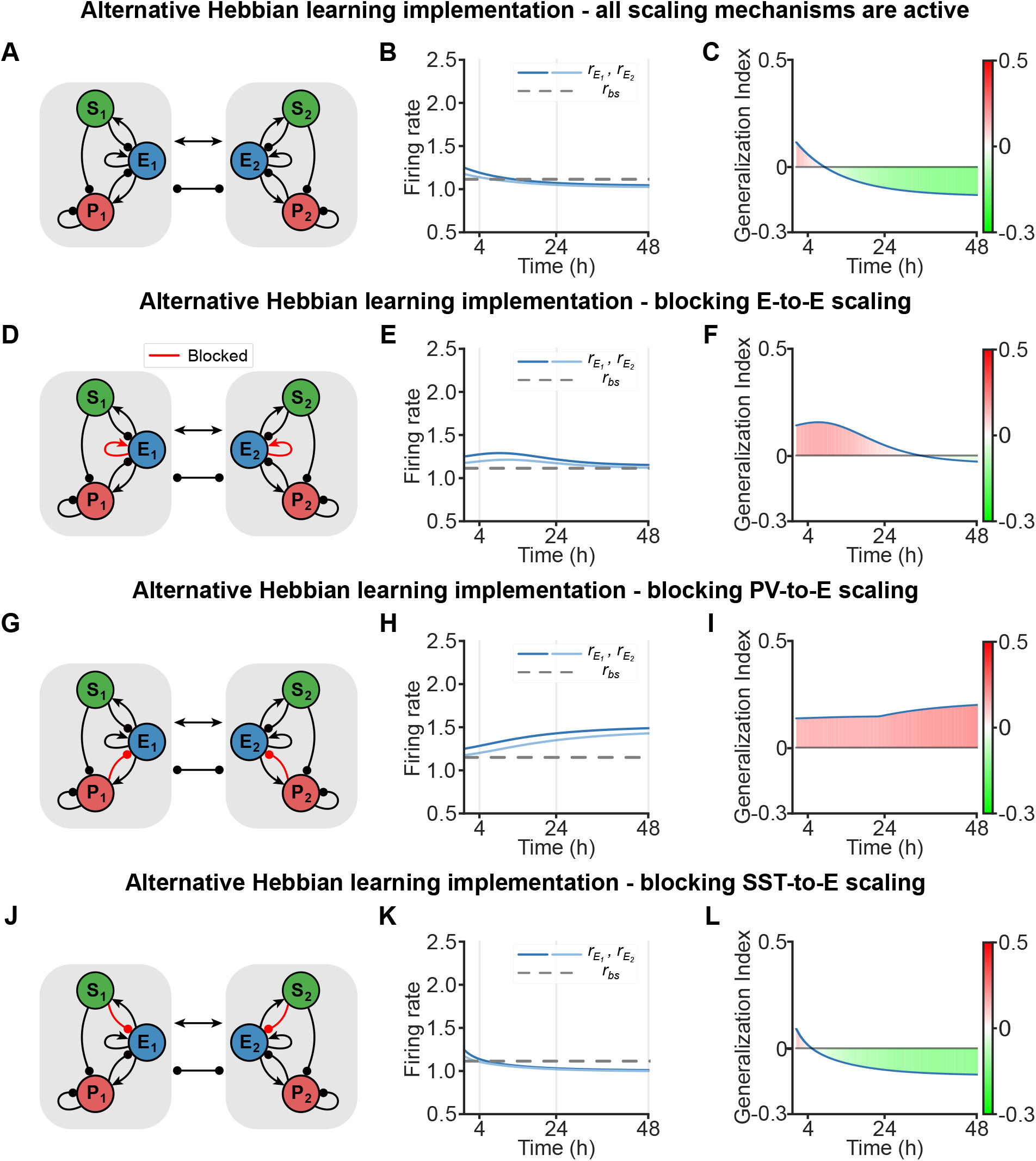
Cell-type-specific synaptic scaling contributions to memory refinement in associative learning with alternative Hebbian learning implementation. **A**. Network schematic when all scaling mechanisms are active, compare to Figure 3F and 4B. **B**. Activity of excitatory population in subnetwork 1 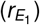 and subnetwork 2 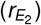 after conditioning. The gray horizontal dashed line represents the baseline activity level measured before conditioning. **C**. Evolution of the Generalization Index (GI) after conditioning. A positive GI indicates memory generalization, whereas a negative GI represents memory specificity. **D - F**. Same as (A - C) but for blocking E-to-E scaling (red connections), compare to Figure 7D - F. **G - I**. Same as (A - C) but for blocking PV-to-E scaling, compare to Figure 7G - I. **J - L**. Same as (A - C) but for blocking SST-to-E scaling, compare to Figure 7J - L.

**Fig. S9.**
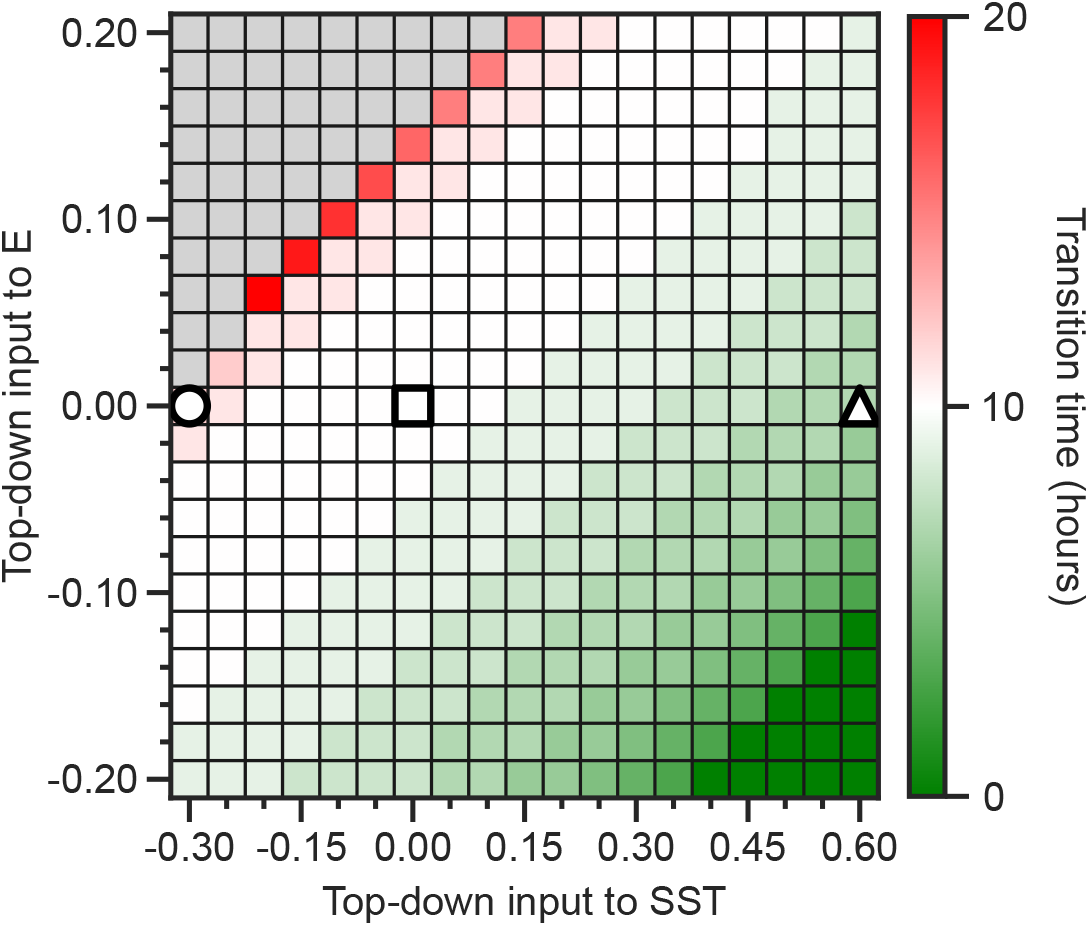
Transition time defined as the time when the memory generalization switches from memory specificity, (e.g. when the Generalization Index switches from positive to negative) as a function of top-down input to excitatory and SST populations. Square, circle and triangle symbols indicate results without top-down inputs shown in Figure 4B, with excitatory top-down inputs to SST shown in Figure 6E, and with inhibitory top-down inputs to SST shown in Figure 6J, respectively.

